# Molecular mechanism of high affinity sugar transport in plants unveiled by structures of glucose/H^+^ symporter STP10

**DOI:** 10.1101/2020.11.05.369397

**Authors:** Laust Bavnhøj, Peter Aasted Paulsen, Jose C. Flores-Canales, Birgit Schiøtt, Bjørn Panyella Pedersen

## Abstract

Sugars are essential sources of energy and carbon, and also function as key signaling molecules in plants. Sugar Transport Proteins (STP) are proton-coupled symporters, solely responsible for uptake of glucose from the apoplastic compartment into cells in all plant tissues. They are integral to organ development in symplastically isolated tissues such as seeds, pollen and fruit. Additionally, STPs play a significant role in plant responses to both environmental stressors such as dehydration, and prevalent fungal infections like rust and mildew. Here, we present two high-resolution crystal structures of the outward-occluded and inward-open conformations of *Arabidopsis thaliana* STP10 with glucose and protons bound. The two structures describe key states in the STP transport cycle. Together with *in vivo* biochemical analysis and Molecular Dynamics simulations they pinpoint structural elements that explain how STPs exhibit high affinity for sugar binding on the extracellular side and how it is considerably lowered on the intracellular side to facilitate substrate release. These structural elements, conserved in all STPs across plant species, clarify the basis of proton-to-glucose coupling, essential for symport. The results advance our understanding of a key molecular mechanism behind plant organ development, and sets the stage for novel bioengineering strategies in crops that could target seeds, fruits and plant resistance to fungal infections.

## Introduction

Correct plant development requires the ability to sense the carbon level of the entire organism. Central to this is a two-step process after unloading of sucrose from the phloem; apoplastic sucrose is enzymatically hydrolyzed to glucose and fructose, then imported into sink tissues by STPs^1,2^. This tightly regulated process is the main driver of monosaccharide uptake, and is essential for correct organ development such as pollen and seed development^3,4^. Besides organ development, STPs function in a wide range of other physiological processes. In guard cells, glucose import by STPs provide carbon sources for starch accumulation and light-induced stomatal opening that is essential for plant growth^5^. STPs play a role in senescence, programmed cell death and participate in the recycling of sugars derived from cell wall degradation^6–8^. STPs and related proteins are implicated in a range of physiological plant responses to environmental stressors, including osmoregulation, salt tolerance, dehydration response and cold response^9^. Sugar uptake by wound and pathogen induced STPs is a central plant immunity strategy; by keeping the apoplast free of sugars, apoplastically growing pathogens like *Pseudomonas syringae* are nutritionally deprived^10–17^. Biotrophic pathogens like the agriculturally important fungi rust and mildew exploit this plant defense mechanism using specialized cell wall penetrating structures called haustoria^18–20^.

The STPs are prominent members of the Sugar Porter (SP) family (also called the MST(-like) family in plants)^21,22^. STPs have a broad pH optimum and display significantly higher sugar affinity (up to 1000x fold) compared to known SP members from other kingdoms^8,23–25^. *Arabidopsis thaliana* STP10 is a canonical STP found in primordia of lateral roots and in pollen tubes. It is a proton driven symporter with a broad pH optimum and with a low μM range affinity for glucose. It also has the ability to transport galactose and mannose^25^. We recently published the structure of STP10 which shows that STPs have a Major Facilitator fold with 12 transmembrane helices constituting two domains, the N domain (helices M1-M6) and the C domain (helices M7-M12), connected by a cytosolic helical bundle domain (IC1-IC5)^26^ (Fig. 1a). On the apoplastic side, a Lid domain (L1-L3) links the N domain to the C domain by a disulfide bridge. Glucose transport could be linked to the protonation of an acidic residue (Asp42) on the M1 helix^26^. The molecular mechanism by which STPs mediate high affinity glucose transport in conjunction with protons is unknown, but has decisive implication on the function of STPs in organ development and plant pathogen defense mechanisms. Here we present a 1.8 Å resolution outward facing structure of STP10, together with a 2.6 Å resolution structure of STP10 in an inward facing conformation, both with glucose bound. The two structures capture two key conformational states in glucose translocation (Fig. 1b). In combination with biochemical characterization and Molecular Dynamics simulations, we show that both structures represent glucose and proton bound states, and address the molecular mechanism of glucose import by STPs.

**Fig. 1.**
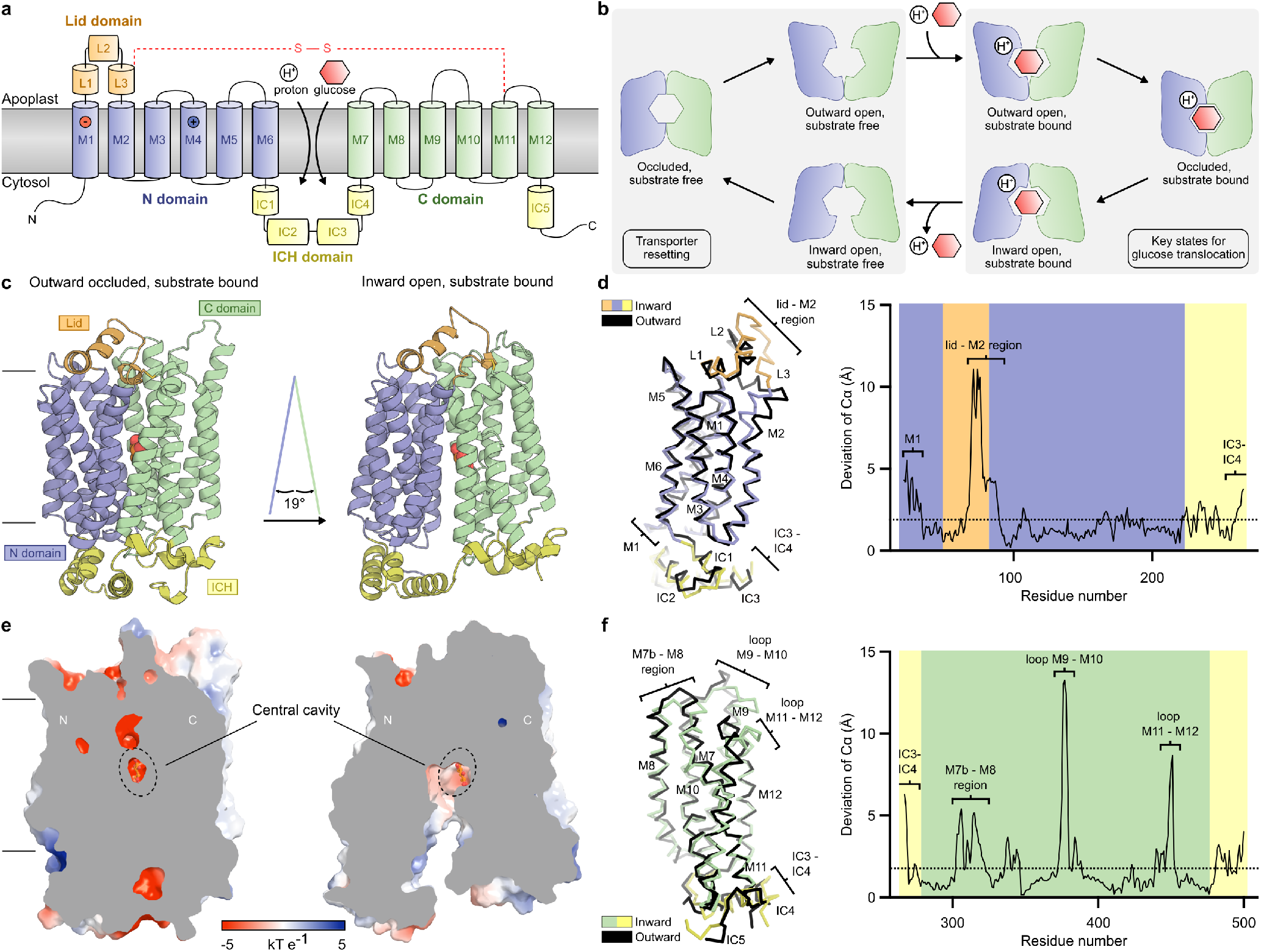
Structures of STP10 in the outward occluded conformation and the inward open conformation. **a)** A simplified diagram of the STP10 topology. Substrate (glucose and proton), the disulfide bridge and position of the proton donor/acceptor pair (M1 and M4) are mapped. **b)** A schematic illustration of the STP10 transport mechanism. Each transport cycle involves binding of substrate in the outward open state and shuttling through a intermediate occluded state to the cytoplasmic side where substrate is released from the inward open state. Returning to a outward open state through a occluded state completes the cycle. **c)** The two solved conformations of STP10 are distinguished by a 19°opening between the N domain (blue) and C domain (green). Glucose (spheres) is buried at the interface between the transmembrane domains above the ICH domain (yellow). Lid domain (orange) is connected to the C domain by a disulfide bridge (sticks). **d)** Left: Superposition of the two conformations using C*α* backbone of N domain residues (21-224 (blue)) including Lid domain (53-80 (orange)) and IC1-3 residues (225-266 (yellow)). Right: Corresponding plot of deviation of the C*α* positions. Significant local conformational changes (>4.0 Å r.m.s.d) between the two structures are observed in M1, lid-M2 region as well as in the IC3-IC4 region as indicated with brackets. Overall r.m.s.d. of the C*α* atoms for the N domain half is 1.93 Å (dotted line). **e)** A slab through the surface electrostatic potential of the STP10 structures. In the outward occluded structure, the glucose is occluded from the intracellular and extracellular side, whereas the glucose is solvent-accessible from the intracellular side in the inward open structure. **f)** Left: Superposition of the two conformations using C*α* backbone of C domain residues (267-500 (green)) including IC4 residues (267-281 (yellow)) and IC5 residues (476-500 (yellow)). Right: Corresponding plot of deviation of the C*α* positions. Significant local conformational changes (>4.0 Å r.m.s.d) between the two structures are observed in IC3-IC4 region, M7b-M8, M9-M10 loop and M11-M12 loop as indicated with brackets. Overall r.m.s.d. of the C*α* atoms for the C domain half is 1.83 Å (dotted line).

## Results and Discussion

### Structures of STP10

We determined the crystal structure of STP10 in two different substrate-bound conformations: outward occluded at 1.8 Å and inward open at 2.6 Å resolution (Extended Data Fig. S2). The new outward occluded structure is overall similar to the previously published structure with an r.m.s.d. (C*α*) of 0.396 Å. However the improved very high resolution of 1.8 Å provides a highly detailed atomic structure of STP10 (residues 21-507, Rfree 21.2%) (Fig. 1c, Extended Data Figs. S3 and S4, Extended Data Table S1). The structure includes a glucose molecule in the central binding pocket with the N and C domains clamped around it (Fig. 1e). Exit from the binding site towards the cytosol is completely blocked and held in place by several strong interactions, including three prominent salt bridges between the N and C domain at the cytosolic interface (Fig. 1 and Extended Data Fig. S5a). The residues that create the salt bridges are strictly conserved in all Sugar Porters, and constitute the canonical MFS and Sugar Porter signature motifs called the A motif and SP motif, respectively (Extended Data Fig. S1)^27^. In other SP proteins, these residues play a central role in stabilizing an outward facing conformation, and we hypothesized that disruption of this salt bridge network should facilitate arrest of STP10 in an inward-facing state^23,28–35^. To test this, we created an STP10 double mutant (E162Q/D344N) and measured *in vivo* activity. The double mutant abolishes transport activity, supporting a conformational arrest of the transporter (Extended Data Fig. S6a), and the double mutant readily crystallized in an inward-open conformation. The structure was refined to 2.6 Å resolution (residues 16-500, Rfree 27.79%) (Fig. 1c, Extended Data Figs. S3 and S4, Extended Data Table S1). Map quality is high except for a part of the Lid domain (residues 64-73) which is poorly defined, indicative of high flexibility (Extended Data Fig. S3b). In the inward open conformation, the N and C domains tightly interact at the apoplastic side forming an inverse V-shaped structure with an open cavity extending 25 Å from the cytosol to the binding site (Fig. 1c,e). In this central binding site, a single glucose molecule is present.

The two obtained conformations represent the two major states in a transport mechanism that cover the translocation of glucose from the apoplastic space to the cytosol (Fig. 1b). Structural alignment between them show intradomain rearrangement of the Lid domain, ICH domain, N domain and C domain during the transition (Fig. 1d,f). The M2-Lid region and helix-loop-helix region of M9-M10 display dramatic rearrangement, while significant differences are also observed in M1, region M7b-M8 and region M11-M12 (Fig. 1d,f). In the inward open conformation, IC1, IC2, and IC5 maintain a well-defined and similar conformation with respect to the N and C domains, but region IC3-IC4 are stretched compared to the outward conformation to allow STP10 to open to the cytosol (Fig. 1c,d,f). The two structures reveal that the domains of STP10 remain rigid in large areas, while exhibiting local rearrangements linked to glucose and proton translocation during the transport cycle. While the Lid domain undergoes large movements between the two conformations the Cys77(Lid)-Cys447(M11) disulfide bridge, which links the Lid and N domain to the C domain, is well defined and clearly visible in both conformations (Fig. 2a).

**Fig. 2.**
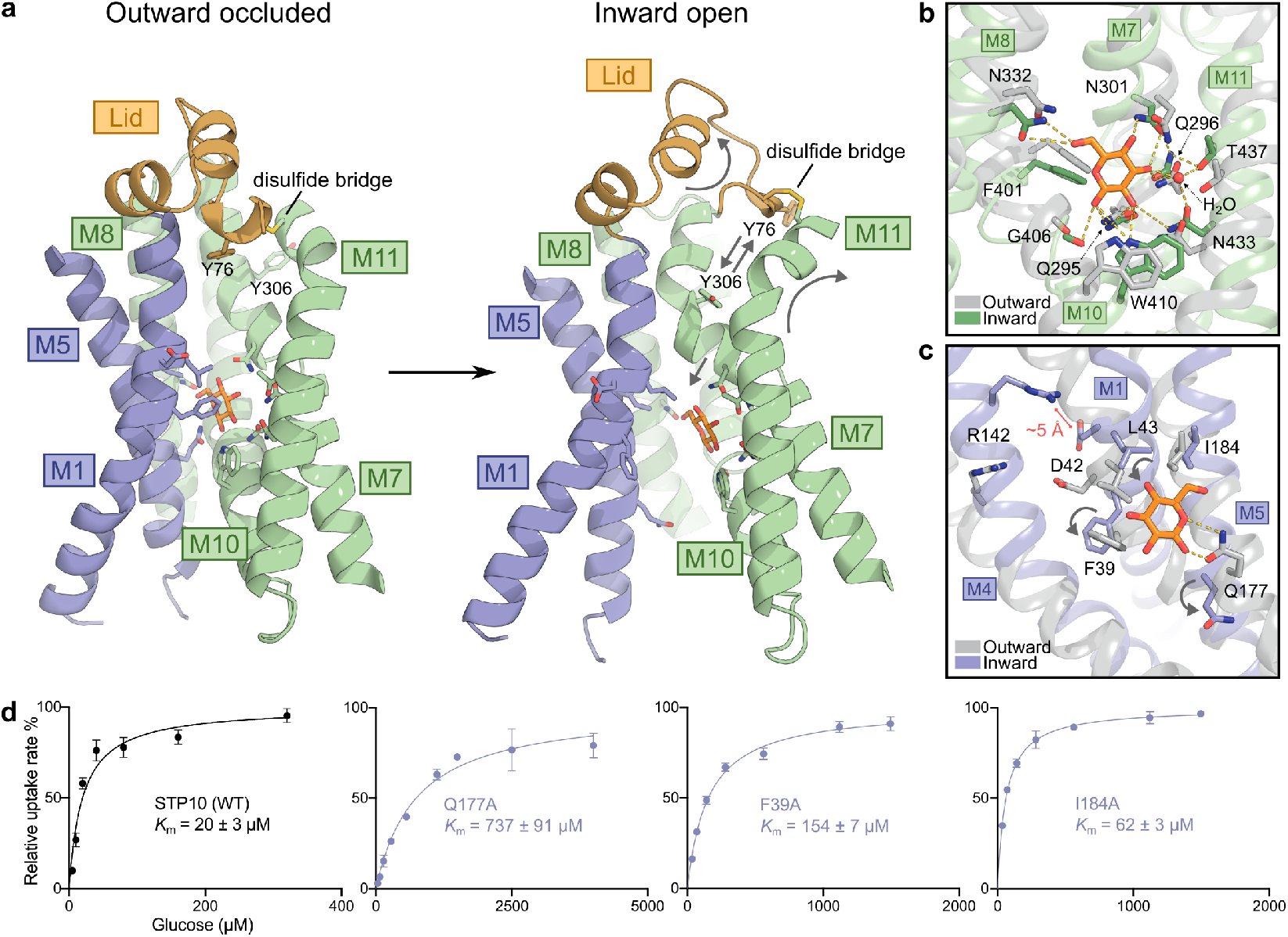
Structures of STP10 with bound glucose and uptake. **a)** Structural changes of the central cavity following outward occluded to inward open transition. The central binding site of STP10 is lined by conserved residues from M1 and M5 of the N domain and M7, M8, M10 and M11 from the C domain. Arrows indicate major changes. **b)** The glucose binding site towards the C domain in the inward (green) and the outward (grey) structure superposed on glucose. Yellow dashes indicate hydrogen bonds. **c)** The binding site towards the N domain and the proton donor/acceptor pair in inward (blue) and outward (grey) structure superposed on glucose. Grey arrows indicate major changes. **d)** Michaelis-Menten analysis of glucose uptake of WT and N domain mutants Q177A, F39A and I184A. Data represents mean ±SD of three or more replicate experiments.

In the inward open structure, the glucose density matches the density of the glucose molecule bound in the outward occluded state of STP10, there is no difference in binding pose of the sugar (Fig. 2a-c, Extended Data Fig. S4). In both states, interactions between protein and glucose are primarily mediated by C domain residues from M7, M8, M10 and M11 that make multiple polar contacts to the glucose. No side chain rearrangements are observed during transition (Fig. 2a,b). From the N domain, a few specific interactions with M1 and M5 dominate (Fig. 2a). The inward open substrate bound structure shows that during transition, Phe39(M1) and Gln177(M5) move from close contact with glucose (distances of 3.9 Å and 2.6 Å), to more than 8 Å and 10 Å away from the glucose, while Leu43(M1) and Ile184(M5) maintain close contact (Fig. 2c). The displacement of Phe39 and Gln177 lower the affinity towards the substrate significantly, supported by uptake assays. The F39A and Q177A mutants lead to an almost 8-fold (*K*_m_ 154 μM) and 37-fold (*K*_m_ 737 μM) reduction in affinity, compared to STP10 WT (*K*_m_ 20 μM) (Fig. 2d). In comparison, the I184A mutant reduced affinity by 3-fold (Fig. 2d).

### The proton site is protonated in both substrate-bound conformations

The proton site, constituted by proton donor/acceptor pair Asp42(M1) and Arg142(M4) is essential for the transport ability of STP10^26^. To elucidate the mechanism of proton driven symport, a cornerstone condition is to establish the protonation state of the solved substrate bound structures. In the outward occluded conformation of STP10, crystallized at pH 4.0, the carboxyl-group oxygen atoms of Asp42 and the guanidine group nitrogen atoms of Arg142 are 4.7 Å apart and stabilized by an acetate ion from the crystallization cocktail (Figs. 2c and 3a). In the inward open substrate-bound state, crystallized at pH 9.0, this distance is maintained at 5.3 Å (Figs. 2c and 3b). This distance reflects the protonation state, and suggests that Asp42 is neutralized by a proton, as Asp42 is expected to move closer to form a salt bridge with Arg142 in its negatively charged state^26,36^. To test this hypothesis, we used molecular dynamics (MD) simulations. Ten independent repeats for the outward and inward state with Asp42 either neutral or charged were carried out over approximately 2 μs (accumulated simulation time 80 μs). When Asp42 is neutral in both outward occluded and inward conformations, Asp42 and Arg142 maintain a broad distance distribution with a median distance of 5-7 Å (Fig. 3c,d). In particular, in the outward conformation the large distribution reflects the flexibility between the M1 and M4 helices. With a negatively charged state of Asp42 in both conformations conversely, the distance is consistently reduced with a extremely narrow distance distribution centered around a ~2.8 Å salt bridge interaction of Asp42-Arg142 that is formed rapidly and maintained throughout all simulations (Fig. 3c,d). We followed this up by calculating the p*K*_a_ of Asp42 using the free energy perturbation (FEP) method and an ensemble of five independent calculations (Extended Data Table S2). The results predict that Asp42 has a p*K*_a_ of 8.7 (+/- 0.3) in the outward state, and a p*K*_a_ of 14 (+/- 1) in the inward state. A continuum electrostatic calculation on the crystal structures predicts Asp42 p*K*_a_ values of 6.8 and 8.8 for the outward and inward states, respectively, showing a similar trend as the FEP results (Extended Data Figs. S8a,b). We conclude that the aspartate of the proton-site is neutralized by a proton in both conformations.

**Fig. 3.**
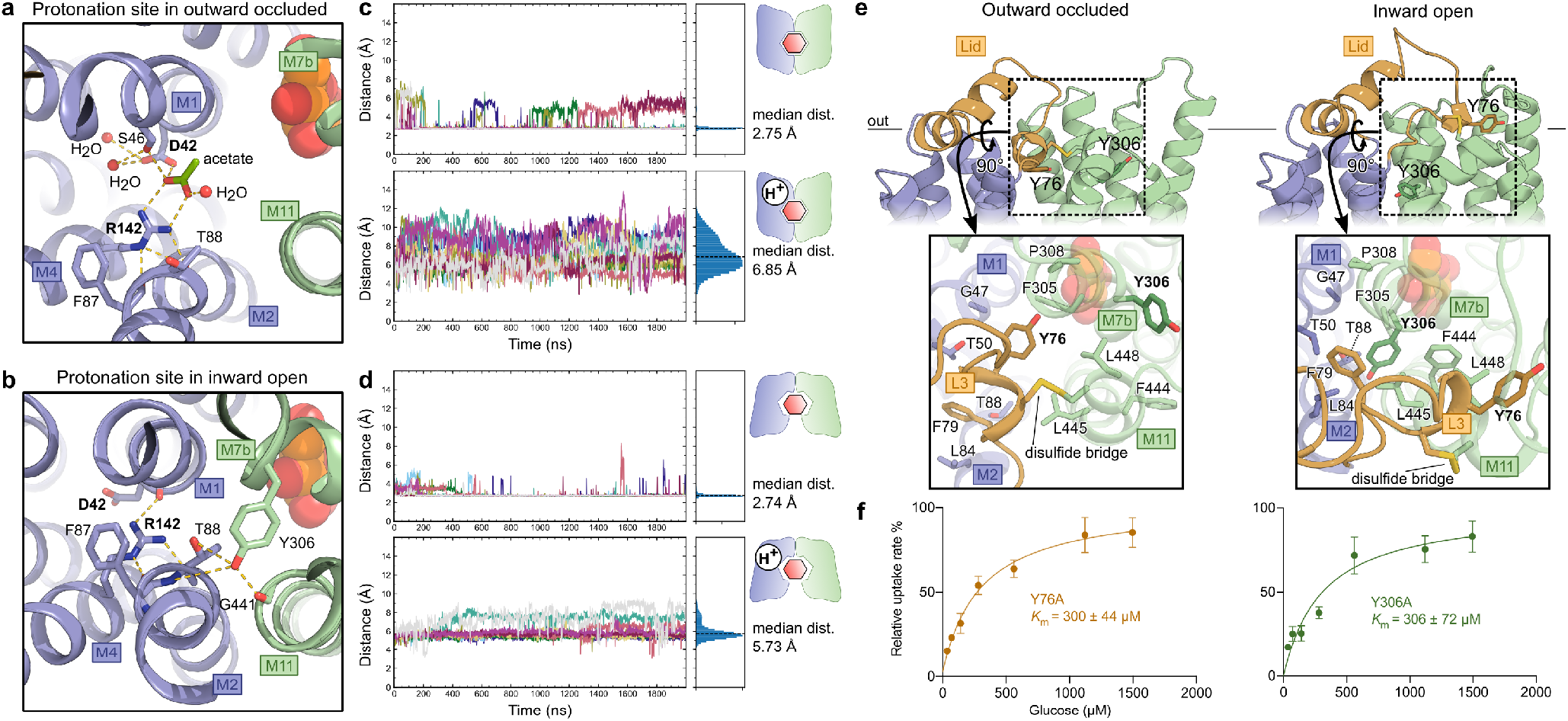
Molecular dynamics simulations of the protonation site and exofacial gating of STP10 states. **a)** The proton-binding site in the outward-facing STP10 structure. Yellow dashes indicate hydrogen bonds (<3.5 Å). **b)** The proton-binding site in the inward-facing STP10 structure. Yellow dashes indicate hydrogen bonds (<3.5 Å). **c)** Distance measurements between Asp42 and Arg142 of the outward-facing state throughout 2 μs MD simulations measured in ten independent repeats, shown in different trace colors. The simulations were carried out for the outward-facing state of STP10 with Asp42 either charged (top panel) or neutral (bottom panel). The median distance between Asp42-Arg142 in the charged state was calculated to 2.76 Å while being 6.85 Å in the neutral state. The distance range for the top panel is [2.46 Å, 8.95 Å] and [2.59 Å, 14.92 Å] for the bottom panel. All distance traces correspond to the minimum distance between the oxygen atoms of the carboxyl group of Asp42 and the nitrogen atoms of the guanidine group of Arg142. **d)** Distance measurements between Asp42 and Arg142 of the inward-facing state throughout 2 μs MD simulations measured in ten independent repeats. The simulations were carried out for the inward-facing state of STP10 with Asp42 either charged (top panel) or neutral (bottom panel). The median distance between Asp42-Arg142 in the charged state was calculated to 2.74 Å while being 5.71 Å in the neutral state. The distance range for the top panel is [2.45 Å, 9.53 Å] and [2.62 Å, 10.92 Å] for the bottom panel. **e)** Close-up views of hydrophobic interactions that enclose the central cavity from the extracellular side in the outward occluded (left) and the inward open (right) structures. **f)** Michaelis-Menten analysis of glucose uptake of Y76A and Y306A. Data represents mean ±SD of three or more replicate experiments.

### Mechanism of occlusion links proton site to glucose site via Lid domain

Next we sought to explain how the proton site is linked to full occlusion of the central glucose binding site during the transition from the outward to the inward facing conformation.

In the outward occluded conformation, the side chain of Arg142 is in contact with the backbones of conserved residues Phe87(M2) and Thr88(M2) (Fig. 3a). In the inward open state, Arg142 reorient to form polar contacts with the backbone of Asp42, but maintains the backbone interaction to Phe87 and Thr88 (Fig. 3b). This locks the movements of the flexible side chain of Arg142 to the M2 helix, and links proton site changes to M2. In the transition to inward-facing, this pulls the M2 helix towards the part of central cavity facing the exofacial side of the protein. This movement enables M2 and M11 to make polar interactions resulting in a pronounced kink in M11 of the inward facing conformation (Fig. 2a). Mutagenesis of Phe87 and Thr88 abolishes transport (Extended Data Fig. S6a).

In the outward occluded conformation, the Lid domain is clamped down on top of the protein with its hydrophobic L3 helix embedded between the extracellular region of the N and C domain (Figs. 1a and 2a). In the inward open conformation, the induced kink of M11 created by the proton site drives the Lid to disengage from the rest of the protein, and the L3 region is exposed to the extracellular space (Figs. 1a and 2a). The high b-factors of the L2-L3 region in the inward open structure suggest high flexibility in this conformation (Extended Data Fig. S2c). The opening of the Lid domain is linked by the Cys77(L3)-Cys447(M11) disulfide bridge to the movements of M11. A bend at Pro308(M7b) is orchestrated by this event, that allows hydrophobic residues of M7b to occupy this space to complete the closure of the extracellular entrance to the glucose site. The introduced kink in M7 generates full closure towards the extracellular side, and is enabled by the proline side chain that breaks the alphahelical hydrogen-bonding pattern. Mutating Pro308 led to a 4-fold reduction in affinity (Extended Data Fig. S6b). Overall, these movements link the proton site to the complete occlusion of the glucose binding site to provide isolation from the extracellular space.

The transition leads to a remarkable mimetic residue-swap in the protein: In the outward conformation, the conserved Tyr76(L3) forms a network of hydrophobic interactions to conserved residues which isolate the proton/donor pair and the saturated binding site from the extracellular space (Fig. 3e). During the transition and the release of the Lid, these interactions are replaced by a reshuffling and new interactions in the same network to Tyr306(M7b) (Fig. 3e). During the transition, the hydrophobic network thus acts as a dynamic outer gate; Tyr76(L3) is rearranged in the Lid domain by moving 19 Å, while Tyr306 from M7b takes its place to control access to the binding site during the transport cycle (Fig. 2a). The mutation of either of the tyrosines (Y76A and Y306A) led to significant impairment of transport activity with an almost identical 15-fold decrease in affinity (*K*_m_ ~300 μM) (Fig 3f). The F79A mutant was used as a negative control and retained affinity comparable to wild-type STP10 (*K*_m_ 29 μM) (Extended Data Fig. S6c).

### A transient chloride site exist at the endofacial side of STP10

In the inward open conformation of STP10, a strong spherical density is present between the SP motif and the A motif of the N domain (Fig. 4a). We replaced Cl^-^ with its chemical congener Br^-^ in our crystallization experiments, which gave rise to a single strong anomalous peak (8.72 sigma) at the position of this spherical density, allowing us to unambiguously identify this peak as a Cl^-^ ion (Fig. 4b,c). In contrast, the outward occluded structure does not contain this peak. Instead a fully conserved aspartate (D225) of the SP motif takes its place and interacts directly with the A motif residues, creating an ‘SP-A network’ that is expected to stabilize the outward conformation (Fig. 4d, Extended Data Fig. S5b). It is noteworthy that the mutant D225N exhibits reduced glucose uptake due to a 5-fold decrease in binding affinity (*K*_m_ 101 μM) (Fig. 4e).

**Fig. 4.**
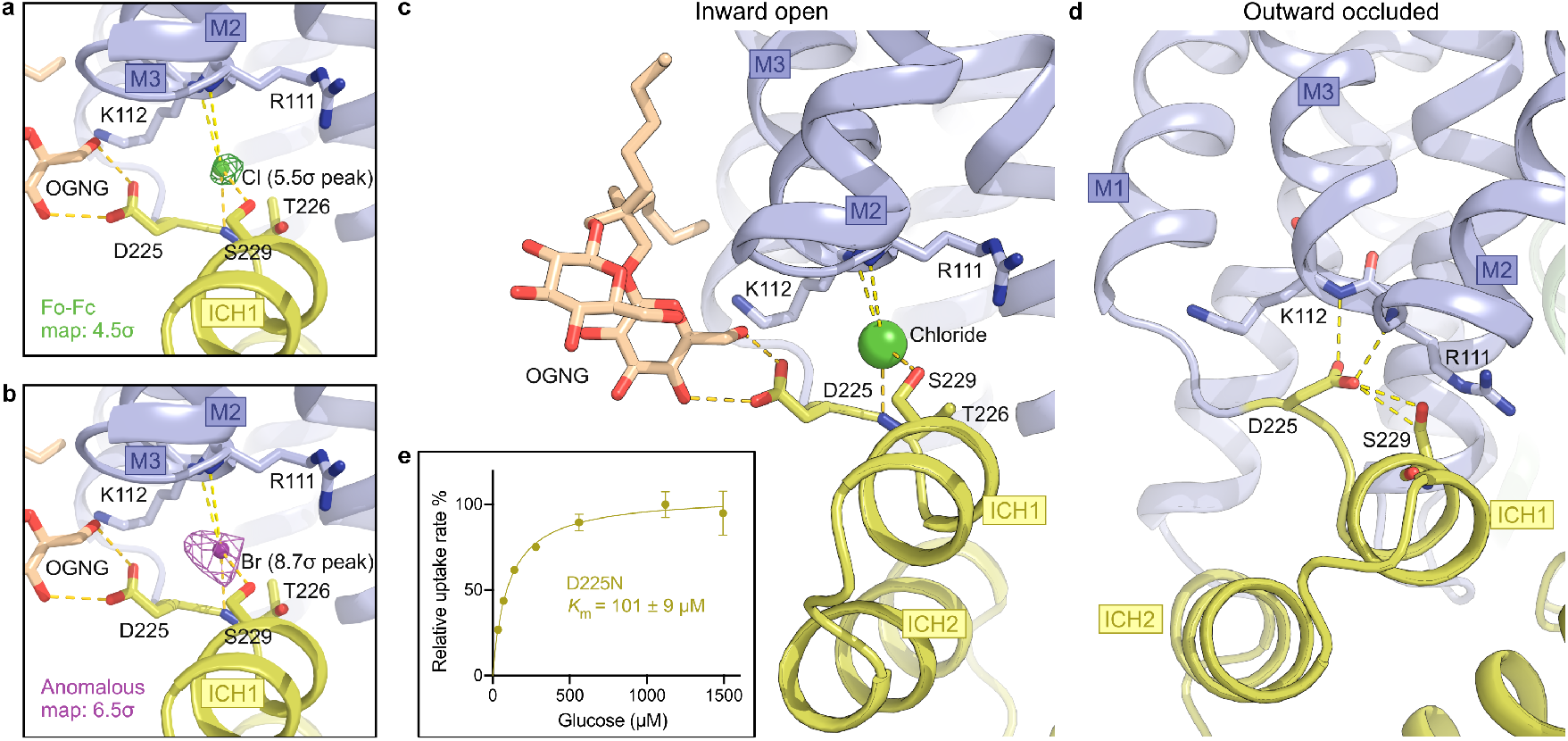
Intracellular molecules target a endofacial regulation site to stabilize the inward-open conformation. **a)** The omit Fo-Fc electron density for the positive density peak contoured in green mesh at 4.5 sigma in the inward-open conformation that was identified as a chloride ion. **b)** The anomalous signal for bromide, shown in magenta mesh contoured at 6.5 sigma. **c)** Intracellular detergent (OGNG) is coordinated to Asp225 in the inward-open conformation. A chloride ion (shown as sphere) neutralizes the A-motif. Selected residues are shown as sticks and hydrogen bonds are represented by yellow dashes (<3.5 Å). **d)** N domain SP-A network in the outward-occluded conformation. Selected residues are shown as sticks and hydrogen bonds are represented by yellow dashes (<3.5 Å). **e)** Michaelis-Menten fit to glucose titration of the D225N at pH 5.0. Data represents mean ± SD of three or more replicate experiments.

Interestingly, in the inward open structure, the Asp225 that is replaced by the chloride, points away from the A motif and interacts with a glucose headgroup of a detergent molecule (Extended Data Fig. S4b and Fig. 4c). It has been shown that direct interaction between lipid headgroups and the conserved cytoplasmic network affects the transition between conformational states in MFS members^35,37^, and while this observed interaction could be an artifact of the crystallization condition, our work support a tentative hypothesis where Asp225 of the SP motif can interact with intracellular glucose or lipid headgroups, modulating transport by a stabilization of the inward open state.

Another endofacial location also warrants our attention. Two cysteines, Cys288(M7) and Cys417(M10), at the intracellular interface of STP10 have thiol groups ~3.5 Å apart from each other in both conformations of STP10 (Extended Data Fig. S4a). We do not observe any disulfide bridge, but the positioning is so striking that we speculate this may function as an intracellular cysteine-based redox regulation site. C288A and C417A mutants display a 3-fold decrease in affinity (*K*_m_ ~60 μM) suggesting an equivalent effect of mutating these residues (Extended Data Fig. S6d,e), but possible involvement in STP10 regulation will require further work. Intracellular cysteine-based redox regulation and signaling has been suggested to occur in plants^38,39^. The cysteine pair is conserved only in STP9, STP10 and STP11 and not found in other *A. thaliana* STPs. Sequence alignment shows that the cysteine pair is found in specific STPs across plant species, supporting a conserved model for isoform specific regulation (Extended Data Fig. S1).

### Model of glucose transport by Sugar Transport Proteins

On the basis of our findings, we propose the following comprehensive model for sugar transport by STPs (Fig. 5). In the outward open state, an open Lid domain allows for a water-filled inlet channel for sugar and protons to the binding sites. Following binding of sugar, protonation and neutralization of Asp42 leads to liberation from the sidechain of Arg142 and distortion of the flexible M1b helix with Phe39, Leu43 and M5 with Ile184 and Gln177 towards glucose, locking it in and creating a high affinity binding pocket. The high affinity binding event orchestrated by the protonation of Asp42 induces movements of M7b, M2 and the outward facing region of the M11 helix that connect the C domain to the Lid domain by a disulfide bridge. This facilitates an enclosure of the Lid domain, mediated by the hydrophobic interactions of Tyr76 at the cavity entrance which locks the two transmembrane domains together. In this outward occluded state, the Lid domain isolates the protonation site from the extracellular space. The transition from outward occluded to the inward open state drives the Lid domain away from the entry site of the central cavity by the bending movement of M11 through the disulfide bridge. This transition is governed by Tyr76 and Tyr306 that swap their position to maintain tight hydrophobic interactions between the central helices keeping both the protonation and binding site isolated from the extracellular space. The transition results in movements of N domain residues Leu43, Ile184 and especially Phe39 and Gln177 away from the bound substrate. This creates a transition from a high sugar binding affinity state to a considerably lowered affinity state in this inward open conformation and the sugar is released. In this inward open state, the SP-A network is broken, as Asp225 is flipped away from the A motif, and a chloride ion takes its position.

**Fig. 5.**
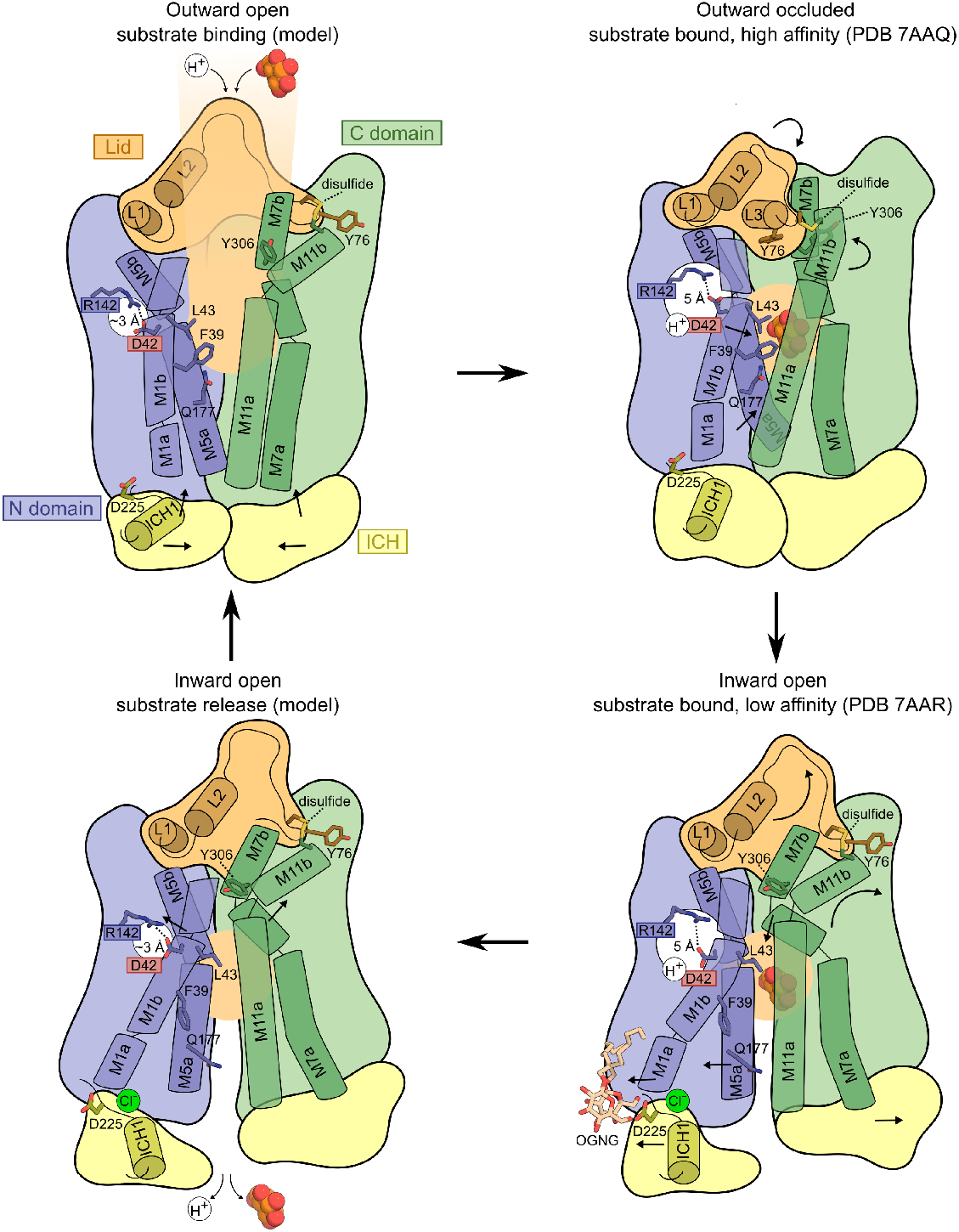
Proposed mechanism of high affinity transport of glucose by STP10. In the outward open conformation (top; left), protons and substrate enter the central binding site. Protonation of Asp42 pushes the flexible M1b towards the sugar binding site, creating a high affinity state that results in enclosure of the Lid domain mediated by M11 movements through the disulfide bride. The Tyr76 residue of the Lid domain ensures that protective enclosure is retained (top; right). A transition to the inward open state result in an opening of the Lid domain bridged to movements of the M11 kink (bottom; right). M7b moves into the central cavity where Tyr306 has switched position with Tyr76 to retain the exofacial gate. Local movements in M1 and M5 results in displacement of substrate binding residues and lowers sugar affinity. Following the transition, the N domain SP motif aspartate flips away from the A motif, replaced by a chloride ion. The lowered affinity and flexibility of M1 (due to neutral Asp42) favor sugar dissociation. Release of the sugar allows Asp42 to close with Arg142 which directs p*K*_a_ changes that favor deprotonation (bottom; left). Deprotonation and the Asp42-Arg142 salt bridge destabilize the inward open state and induce transition towards a more stable outward open state favored by interactions of the intracellular networks of salt bridges and the N domain SP-A network.

In MD simulations of the inward open state, bound glucose can escape to the intracellular side within a few hundred nanoseconds irrespective of the protonation state of Asp42, but a protonated Asp42 appears to favor sugar dissociation (Extended Data Fig. S7a,b). Unloading of glucose results in a displacement of the M1b helix away from the binding site, allowing the neutral Asp42 to move into close proximity of Arg142 which induces proton release. The sugar and consequent proton release reestablishes the interaction between Asp42 and Arg142 thereby destabilizing the inward open state. This shifts the transition equilibrium towards a more stable outward facing state that is favored by the networks of salt bridges and the N domain SP-A network at the intracellular side. The outward open state is then ready to allow substrate and protons to enter the central cavity for another transport cycle.

This transport mechanism is likely broadly conserved within the SP protein family found in all plants. Alongside STPs, the family also includes the ERD6-like, PMT, pGLcT, VGT, TST and INT subfamilies^22^. Members of these subfamilies can play determinant roles in plant development and tolerance to environmental stress. For instance, INT and ERD6-like members are involved in the regulation of cell elongation and in responses to abiotic stress like dehydration, respectively^40,41^. Our work pinpoints crucial structural elements, conserved across both protein subfamilies and plant species, and explain their importance for high sugar affinity transport (Extended Data Fig. S1). Strikingly, single point mutations in these key elements of the transport mechanism of STPs mediate rust and mildew resistance in wheat and barley, and our work provides the foundation for leveraging this knowledge in new bioengineering strategies in crops^12,20^.

In conclusion, we have presented two key state structures of a STP protein together with biochemical and Molecular Dynamics simulations that explain high affinity glucose transport in plants. The continued structural and functional characterization of plant sugar transporters is key not only for a molecular understanding of fundamental physiological processes in all plants, but also from an applied point of view to address future challenges in biotech, agriculture and environmental sciences.

## Acknowledgements

The authors acknowledge beamlines I24 and I04 at the Diamond Light Source and beamline BioMAX at the MAX IV Laboratory, where X-ray data were collected, as well as DESY-PETRA III for crystal screening. This work was supported by funding from the European Research Council (grant agreement No. 637372), the Danish Council for Independent Research (grant agreement No. DFF-4002-00052), the Carlsberg Foundation (CF17-0180), and an AIAS fellowship to B.P.P. Novo Nordisk Foundation (NNF18OC0052988), the Villum Foundation (project number 34326), and the Independent Research Fund Denmark, Natural Sciences (7014-00192B) supported J.C.F.-C. Computations were performed at the Grendel-S cluster of the Centre for Scientific Computing Aarhus (CSC-AA), and made possible by a grant from the Novo Nordisk Foundation (NNF18OC0032608).

## Author contributions

L.B. did crystallization experiments, processed data, and biochemical characterization. P.A.P. did crystallization experiments and processed data. J.C.F.-C. performed molecular dynamics simulations. B.S. supervised the molecular dynamics simulations. B.P.P. supervised the project. L.B. and B.P.P. wrote the paper. All authors commented on the paper.

## Author information

Coordinates and structure factors have been deposited in the Protein Data Bank with the accession numbers 7AAQ (outward) and 7AAR (inward). The authors declare no competing interests. Correspondence and requests for materials should be addressed to B.P.P.

## Methods

### Protein Purification

The gene encoding *Arabidopsis thaliana* STP10 (UniProt: Q9LT15) was introduced into an expression construct based on p423_GAL1 with a C-terminal purification tag containing a thrombin cleavage site and a deca-histidine tag. To obtain the inward open state the mutations E162Q and D344N were introduced using the quickchange site-directed mutagenesis kit (Agilent). Transformed *Saccharomyces cerevisiae* (strain DSY-5) were grown in a culture vessel to high density by fed-batch and harvested after a 22 hour induction using galactose^42^. Harvested cells were washed in cold water, spun down and re-suspended in lysis buffer (100 mM Tris pH 7.5, 600 mM NaCl, 1.2 mM phenylmethylsulphonyl fluoride (PMSF)), followed by lysing using bead beating with 0.5 mm glass beads. The homogenate was centrifuged for 20 minutes at 5,000g, followed by sedimentation of membranes by ultracentrifugation at 200,000*g* for 2 h. Membrane pellets were re-suspended in membrane buffer (50 mM Tris pH 7.5, 500 mM NaCl, 20% glycerol) before being frozen in liquid nitrogen. 9 grams of frozen membranes were thawed and solubilized for 30 minutes in a solubilization buffer (150 mM NaCl, 50 mM Tris pH 7.5, 5% Glycerol, 50 mM D-glucose, 1% n-dodecyl-β-d-maltoside (DDM) and 0.1% Cholesterol hemi succinate (CHS)) in a total volume of 100 ml, after which unsolubilized materials were removed by filtration using a 1.2 μm filter. 20 mM imidazole pH 7.5 was added and the solubilized membranes were loaded on a pre-equilibrated 5 ml Ni-NTA column (GE Healthcare) at 3 ml/minute. After loading, the column was washed with 10 column volumes of W60 buffer (Solubilization buffer with 0.1% DDM and supplemented with 60 mM Imidazole pH 7.5), followed by a 20 column volumes wash with G-buffer (20 mM Mops pH 7.5, 250 mM NaCl, 10% Glycerol, 0.12% Octyl Glucose Neopentyl Glycol (OGNG), 0.012% CHS, 0.5 mM tris(2-carboxyethyl)phosphine (TCEP)). The composition of the G-buffer was optimized through a thermostability assay^43^. The protein was eluted from the column by circulating 5 ml G-buffer supplemented with bovine thrombin and 20 mM Imidazole pH 7.5, at 19°C for ~16 hours. The following day the column was washed with 15 ml of G-Buffer supplemented with 40 mM imidazole. The samples were pooled and concentrated using a spin column (50 kDa cut-off, Vivaspin) to a volume of ~400 μl and injected on a size-exclusion column (Enrich 650, Biorad), pre-equilibrated in G-buffer. For anomalous experiments to confirm the Cl^-^ site, protein was purified using an identical protocol, but exchanging the G-buffer solution MgCl_2_ to MgBr_2_ and NaCl to NaBr.

### Crystallization

#### Outward occluded state

The peak fractions from SEC were concentrated to ~15 mg/ml using a 50 kDa concentrator (Vivaspin). The outward open state of STP10 was crystallized in lipidic cubic phase (LCP). To prepare lipidic cubic phase for crystallization trials, the protein was supplemented with 100 mM D-glucose before mixing with a 80% monoolein (Sigma-Aldrich) 20% cholesterol mixture, in 1:1.5 protein to lipid/cholesterol ratio (w/w) using a syringe lipid mixer. For crystallization, 50 nl of the meso phase was mixed with 1000 nl of crystallization buffer for each condition on glass sandwich plates using a Gryphon robot (ARI). Tiny crystals appeared after one-two days at 20°C. These crystals diffracted to ~3 Å at Diamond Light Source beamline I24. The crystallization conditions were further optimized and the final optimized crystallization screen contained 0.1-0.15 M Ammonium Acetate, 0.1 M Sodium Citrate pH 4.0 and 36-40% PEG400. This gave crystals with a size of approximately 100×40×40 μm. The crystals were collected using dual thickness micromounts (MiTeGen) and immediately flash frozen in liquid nitrogen. These crystals diffracted to better than 2 Å with a few crystals diffracting anisotropic to 1.6 Å. The final datasets were collected at Diamond Light Source beamline I24 using a wavelength of 0.9686 Å.

#### Inward open state

Peak fractions from SEC with 4-5 mg/ml of protein was used directly for crystallographic experiments. Crystals were grown at 20°C by vapor diffusion in 0.6+0.6 μl sitting drops using MRC Maxi Optimization plates (SWISSCI). The crystals appeared using reservoirs containing 0.3 M NaCl, 0.1 M MgCl_2_, 0.1 M Bicine pH 9.0 and 36-43% PEG400. The crystals appeared after one day and grew to a final size of 150×60×60 μm within 14 days. Data were collected at the Diamond Light Source Beamlines I04 and I24. Crystals that had grown for 3 days diffracted to 3-4 Å whereas crystals that had grown for 14 days diffracted to 2.5-3 Å. One data set was collected from a single crystal with diffraction to 2.64 Å using a wavelength of 0.98 Å. For anomalous experiments to confirm the Cl^-^ site, crystals were grown using an identical protocol, but exchanging the reservoir solution MgCl_2_ to MgBr_2_ and NaCl to NaBr, before mixing the 0.6+0.6 μL drops. Anomalous data were collected at the BioMAX beamline at MAX IV Laboratory using a wavelength of 0.9203 Å.

### Data processing

#### Outward occluded state

Datasets were processed and scaled using XDS^44^ in space group P212121 (#19), which suggested the presence of one STP10 monomer in the asymmetric unit (~58% solvent content). Molecular replacement was done using Phaser^45^ and the outward open STP10 (PDB: 6H7D) as the search model. Afterwards the model was optimized further by running phenix.refine^46^. Final refinement in phenix.refine was done with a refinement strategy of individual sites, individual ADP, and group TLS (3 groups), against a maximum likelyhood (ML) target with reflections in the 50-1.8 Å resolution range. The final model yielded a Rwork of 18.74% and Rfree of 21.20% (Extended Data Table 1). MolProbity^47^ evaluation of the Ramachandran plot gave 99.18% in favored regions and 0.0% outliers.

#### Inward open state

Datasets were processed and scaled using XDS^44^ in space group C2221 (#20), which suggested the presence of one STP10 monomer in the asymmetric unit (~64% solvent content). To solve the phase problem, Molecular Replacement (MR) was done using Phaser^45^ and a search model that contained the transmembrane region of the N-domain from the previously published STP10 (PDB: 6H7D). The partial solution was then used as input for a second round of MR, now using the transmembrane region of the C-domain as the search model. Some of the missing loops were then build manually followed by Molecular Dynamics based geometry optimization using MDFF^48^ through Namdinator^49^. After this, the model could be further improved by iterative manual model building in COOT^50^ combined with Rosetta optimization in phenix.rosetta_refine^51^ and refinement using phenix.refine^46^ guided by 2mFo-DFc maps and Feature Enhanced maps^52^ using model phases. Final refinement in phenix.refine was done with a refinement strategy of individual sites, individual ADP, and group TLS (2 groups), against a maximum likelyhood (ML) target with reflections in the 20-2.64 Å resolution range. The final model yielded a Rwork of 24.66% and Rfree of 27.79% (Extended Data Table 1). MolProbity^47^ evaluation of the Ramachandran plot gave 96.27% in favored regions and 0% outliers. Crystals grown with bromide diffracted to 3.2 Å. Data were collected at the wavelength near the bromine K-absorption edge (0.9203 Å) to maximize the anomalous signal, and processed to 3.5 Å in XDS. The anomalous difference Fourier map was calculated based on data from 18 to 6.5 Å resolution and model phases. One single strong anomalous peak (8.72 sigma) was identified, confirming the bromide/chloride site.

#### Data analysis

All structural figures were prepared using PyMOL (The PyMOL Molecular Graphics System, Version 1.5.0.4 (Schrödinger LLC, 2012)). Superposition of the two protein structures was carried out by matching graphs built on the protein’s secondary-structure elements, followed by an iterative three-dimensional alignment of protein backbone C-alpha atoms using superpose^53^. Sequence alignments were constructed with PROMALS3D^54^, followed by manually refining gaps based on the transmembrane regions observed in the STP10 structure and predicted for other sequences using Phobius^55^. Alignment was visualized using ALINE^56^. Surface electrostatic potential was calculated using the APBS Electrostatics^57^ plugin with default settings in PyMOL.

### Yeast uptake assay

For functional characterization, experiments were performed essentially as described by Sauer and Stadler^58^. In brief, the STP10 gene was subcloned into a p426MET25 vector^59^ for constitutive expression and transformed into the *S. cerevisiae* hexose transport deficient strain, EBY-WV4000^60^, using the lithium acetate/PEG method. Transformed cells were plated in synthetic dropout media with 2% maltose and without uracil. Four to five colonies were used to inoculate 50 ml of synthetic dropout media with 2% maltose, without uracil and methionine and grown to an optical density at 600 nm (OD600) of ~1.5. Cells were washed twice with 25 mM NaPO_4_ buffer pH 5.0, and resuspended in the same buffer to an OD600 of 10. For each reaction 20 μl of cells were mixed with 180 μl of 50 mM NaPO_4_ adjusted to pH 5.0. Cells were shaken in a thermomixer at 30 °C and tests were initiated by adding substrate. After timed incubation, yeast cells were collected via vacuum filtration on mixed cellulose ester filters (0.8 μm pore size) and immediately washed three times with an excess of ice-cold distilled water. Incorporation of radioactivity was determined by scintillation counting. For all assays 1 μCi [3 H]-D-glucose (PerkinElmer, USA) was used as the radioactive tracer. For the determination of *K*_m_ values, cells were incubated with [3 H]-D-glucose for 4 min to keep uptake in the linear range. For the *K*_m_ value determination, the data was normalized to the predicted *V*_max_ by fitting the data to Michaelis-Menten kinetics. The experiments were performed at least in triplicate and showed similar results. Data was analyzed with GraphPad Prism 8.

### Molecular dynamics simulations

#### Model building

All-atom models were built using the outward and inward STP10 structures from this study. Non-protein molecules were removed except five interacting water molecules within 6 Å of Asp42 and Arg142 in the outward structure and three water molecules in the inward protein structures (one water molecule within 3 Å of Asp42 and Arg142 and the two other ones buried between Asn188 and Met304 in the transmembrane protein region). Hydrogens were added to the initial outward and inward structural models and the structures were minimized using Maestro 2019v1 (Schrodinger LLC, 2019)). In addition, *in silico* mutations were performed in the inward protein structure to restore the wild type sequence.

#### Protonation states assignment

The resulting protein atom coordinates of the model building process were used to estimate the p*K*_a_ values for all titratable sites using continuum electrostatic calculations following a similar protocol to the H++ server^61^. The sugar molecules were removed from the models for these calculations, since they are not explicitly modeled. Electrostatic calculations were performed with the MEAD 2.2.7 package and the internal protein (*ε_p_*) and water dielectric constant (*ε_w_*) were set to 6 and 80, respectively^62^. Hydrogen atoms were added and the amber FF14SB electrostatic charges and mbondi2 radii set were assigned to the protein models using the tLeap program available in AmberTools18^63,64^. pK_1/2_ values, pH at which a titratable site is 50% ionized, for all titratable sites were calculated using Monte Carlo (MC) calculations implemented in the MCTI program using 1000 full MC and 10000 reduced MC steps^65^. The latter is a statistical mechanics approach to account for the interactions of multiple titratable sites in all possible configurations. We also calculated p*K*_a_ values using the empirical method Propka3.0 as a reference^66^. Results are reported for residues showing large pK_1/2_ shifts in Extended Data Fig. S8a,b) for different values of *ε_p_* ranging from 4, 6 to 10. pK_1/2_ shifts decrease as the protein dielectric constant increases, which accounts for the solvent screening effect. Here, we used *ε_p_* = 4 accounting for a relatively stable protein structure embedded in a low dielectric environment such as the membrane. STP10 functions over a pH range between 5 to 7, to setup the ionization states in our MD simulations we used a mid pH value of 6. At pH 6, Glu354’s pK_1/2_ was higher than the desired pH as well as it is partially exposed to the lipid environment. Since our continuum electrostatic calculations do not consider the low dielectric lipid bilayer, we provide a first-order approximation of the change of p*K*_a_ (Δp*K*_a_) of charging Glu354 in the membrane environment. For this we used APBSmem, a software specifically designed to estimate this additional change of p*K*_a_, with *ε_p_* = 4 and *ε_membrane_* = 2, and other recommended parameters (see reference for details)^67^. Δp*K*_a_ is estimated around +15 for both STP10 structures, thus resulting on pK_1/2_ + Δp*K*_a_ > 22 well above pH 6. Therefore, Glu354 is set to be neutral in both outward and inward structures. Furthermore, predicted Asp42’s pK_1/2_ values for both STP10 crystal structures are higher than pH 6, and the additional change Δp*K*_a_ when charging Asp42 in the membrane is −0.1 by APBSmem for both protein structures. Thus, our continuum electrostatic calculations for Asp42 predict pK_1/2_ of 6.8 and 11 for the outward and inward crystal structures, respectively. Note that both the predicted pK_1/2_ difference of Asp42 in the two crystal structures and the buried location of Asp42 inside the protein indicate a possible role of ionization changes of Asp42 in triggering protein conformational changes. Hence, protonation changes of Asp42 will be studied through extensive MD simulations and free-energy perturbation calculations as described in the following subsections. Three residues Glu53, Glu54 and Glu64 located in the extracellular loops L1 and L2 have predicted pK_1/2_ values higher/lower than pH 6, but these residues were set to their default ionization states because of their solvent exposure and to facilitate comparison between different MD simulations focused on changes of protonation of Asp42. The protein contains six histidines, which were set to neutral states (epsilon microstate) for both protein states since they are also exposed to the solvent and to keep the same ionization configuration in all MD simulations. All other titrable sites were set to their standard states.

#### System preparation

Protein structure orientation in a lipid membrane was estimated with the PPM server^68^. Using the predicted orientations, protein-membrane systems were built using the CHARMM-GUI Membrane Builder^69,70^. Different system models for the outward and inward states were built with Asp42 charged or neutral. Terminal chains were capped using acetyl (ACE) and N-methyl amide (NME) groups. A disulfide bridge was built between Cys77 and Cys449. The protein was inserted in a membrane containing approximately 222 POPC lipids and solvated by 19000 to 20000 TIP3P waters providing 18 Å separation to the box edges above and below the protein. The approximate box size for all systems is 96 Å x 96 Å x 110 Å. CHARMM36m force field was used to model the protein, CHARMM36 for the lipids and for the *β*-D-glucose substrate^71–73^. The system charge was neutralized by Cl^-^ counterions.

#### Molecular Dynamics simulations protocols

MD simulations were carried out using the GPU accelerated version of PMEMD18 MD module of the Amber suite of programs^74,75^. The system was first minimized by 2500 steps of steepest-descent followed by 2500 steps of conjugate gradient. Positional restraints where placed in all heavy-atoms of the protein and substrate with a force constant of 100 kcal/(mol Å^2^). The system was then equilibrated by a five-stage equilibration process. The first stage was an NVT (constant number of particles, volume and temperature ensemble), equilibration with 125 ps and time step of 1 fs with a force constant of 10 kcal/(mol Å^2^). This was followed by semiisotropic NPT (constant number of particles, volume and pressure ensemble), equilibration over 250 ps and a force constant of 2.5 kcal/(mol Å^2^). The third stage had a force constant of 1 kcal/(mol Å^2^) over 500 ps with time step of 2 fs. For the following stages, the time step was kept at 2 fs. The fourth stage was 500 ps with 0.5 kcal/(mol Å^2^). The final stage was carried out over 20.5 ns with a weak restrain of 0.1 kcal/(mol Å^2^). For production MD simulations the protein and substrate heavy-atom restraints were released and approximately 2000 ns were performed for each independent repeat. The following parameters were set for all MD simulations reported here. Covalent bonds formed by hydrogen atoms were constrained using the SHAKE algorithm. A nonbonded cutoff of 12 Å was used with a force-based switching cutoff of 10 Å for the van der Waals interactions. Long range electrostatic interactions were calculated using the Particle Mesh Ewald. The system was simulated with a semiisotropic NPT conditions on the membrane plane. The target pressure of 1.0 bar was regulated by a Monte-Carlo barostat and the temperature of 310 K was controlled by a Langevin thermostat with a friction coefficient of 1 psÅ^-1^. The ig parameter of PMEMD was set to −1 to generate a random seed for the pseudo-random generators in all reported MD simulations.

#### Free energy perturbation calculations

The p*K*_a_ of Asp42 was calculated using the following equations:

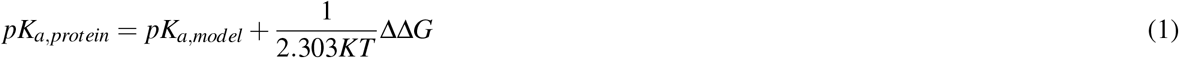

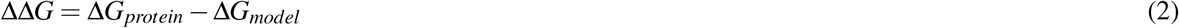

Where ΔΔ*G* is the difference between the free energy of charging Asp42 in the protein environment embedded in a solvated lipid bilayer (Δ*G*_*protein*_) and the free energy of charging Asp in a small dipeptide model in water (Δ*G_model_*). p*K*_a_*model*__ = 4 is the experimentally determined p*K*_a_ of Asp in solution and p*K*_a_, protein is the calculated p*K*_a_ of Asp in a protein environment. *K* is the Boltzmann constant and *T* = 310 K.

Free energy perturbation (FEP) method^76^ was used to calculate Δ*G_protein_* for Asp42 in both outward and inward STP10 structures and Δ*G_model_* for Asp in a dipeptide model solvated in explicit water. FEP is a statistical mechanics method that allows for a gradual transformation of the system from an initial state (neutral Asp) to a new one (charged Asp). In FEP calculations a perturbated Hamiltonian U(X, *λ*) is coupled to a non-physical parameter *λ* as follows:

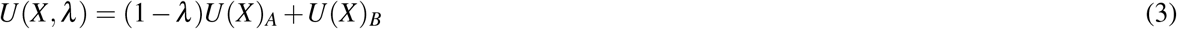

Where *U*(*X*)_*A*_ and *U*(*X*)_*B*_ are the Hamiltonians of the initial and final states *A* and *B* of the system and *X* are the atom coordinates. In the case of charging Asp42 we used the following equation to modify the partial charges of the side chain:

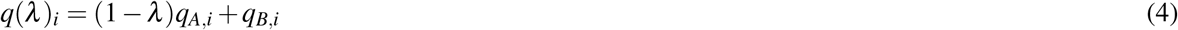

Where *q_i_* is the partial charge of atom *i* in the side chain of Asp42 for states *A* (neutral Asp) and *B* (charged Asp). To prepare the FEP MD simulations, the protein structures were embedded in a POPC lipid bilayer as described in the system preparation. A small peptide of sequence ACE-Asp-NME was built to model aspartate in an unfolded state, it was solvated in a cubic box with a minimum 12 Å distance between the peptide and the simulation box edges. The partial charges of Asp42’s side chain were linearly scaled in 12 states with neutral and charged Asp42 at the initial and final states, respectively^77^. The topology file for each state was edited using Parmed (http://github.com/ParmEd/ParmEd). Hamiltonian replica exchange molecular dynamics (HREMD) was used to improve the convergence of FEP calculations with 500000 exchange trials every 50 MD steps^78^. Note that HREMD implementation in the GPU version of PMEMD18 only supports NVT MD simulations. This required a distinct equilibration process such that all stages have the same box dimensions. An intermediate stage near to the midpoint of the alchemical transformation was first equilibrated using the first four steps of the equilibration protocol described above. The fifth step was carried out over 3 ns followed by a final equilibration process of 5 ns with no restraints. The final NPT equilibrated box was replicated in all intermediate stages and further equilibrated using the corresponding topology file through 5 ns of NVT equilibration with no restraints. FEP calculations were repeated with a neutral form of Arg142, i.e. set to a tautomer (RN2) of the neutral guanidinium side chain, or without a substrate.

#### MD analysis

MD simulation trajectories were analyzed using cpptraj 4.14.0 and MDtraj 1.9.3 software packages^79,80^. VMD 1.9.3 program was used to visualize MD trajectories and calculate surface accessible surface area^81^.

---

**Extended Data Fig. S1.**
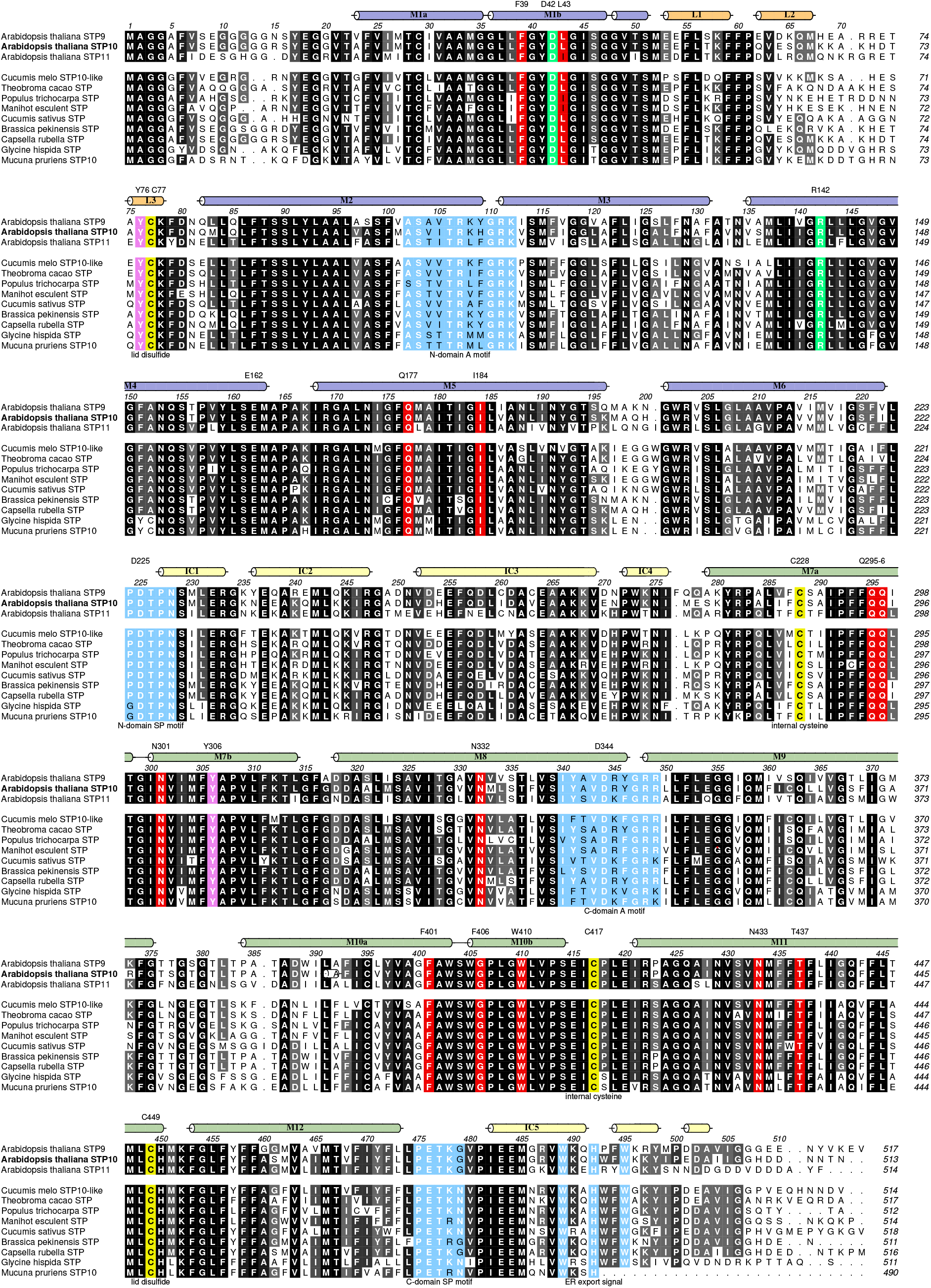
Multiple sequence alignment of the *A. thaliana* Sugar Transport Family STP9, STP10, STP11 with other plant STPs included. Alignment between *A. thaliana* STP9 (accession number Q9SX48), *A. thaliana* STP10 (accession number Q9LT15), *A. thaliana* STP11 (accession number Q9FMX3), *Cucumis melo* cmSTP10-like (accession number A0A5A7SS92), *Theobroma cacao* tcSTP (accession number A0A061E224), *Populus trichocarpa* ptSTP (accession number B9H5Q5), *Manihot esculent* meSTP (accession number A0A2C9V070), *Cucumis sativus* csSTP (accession number A0A0A0LHS6), *Brassica pekinensis* bpSTP (accession number M4FAX8), *Capsella rubella* crSTP (accession number R0I4Q9), *Glycine hispida* ghSTP (accession number I1LF83) and *Mucuna pruriens* mpSTP10 (accession number A0A371FNF1). Conserved residues are highlighted with gray-scale, where black is perfectly conserved. Colored tubes represent *α*-helices found in the N domain (blue), Lid domain (orange), ICH domain (pale yellow) and C domain (green). Key residues are numbered above the *α*-helix markings. Residues highlighted in red participate in sugar binding. The proton donor/acceptor pair is highlighted in green. The cysteines forming the disulfide bridge between Lid domain and C domain as well as the cysteines at the intracellular interface are highlighted in yellow. The tyrosines involved in exofacial gating are highlighted in magenta. Conserved motifs are highlighted in light blue.

**Extended Data Fig. S2.**
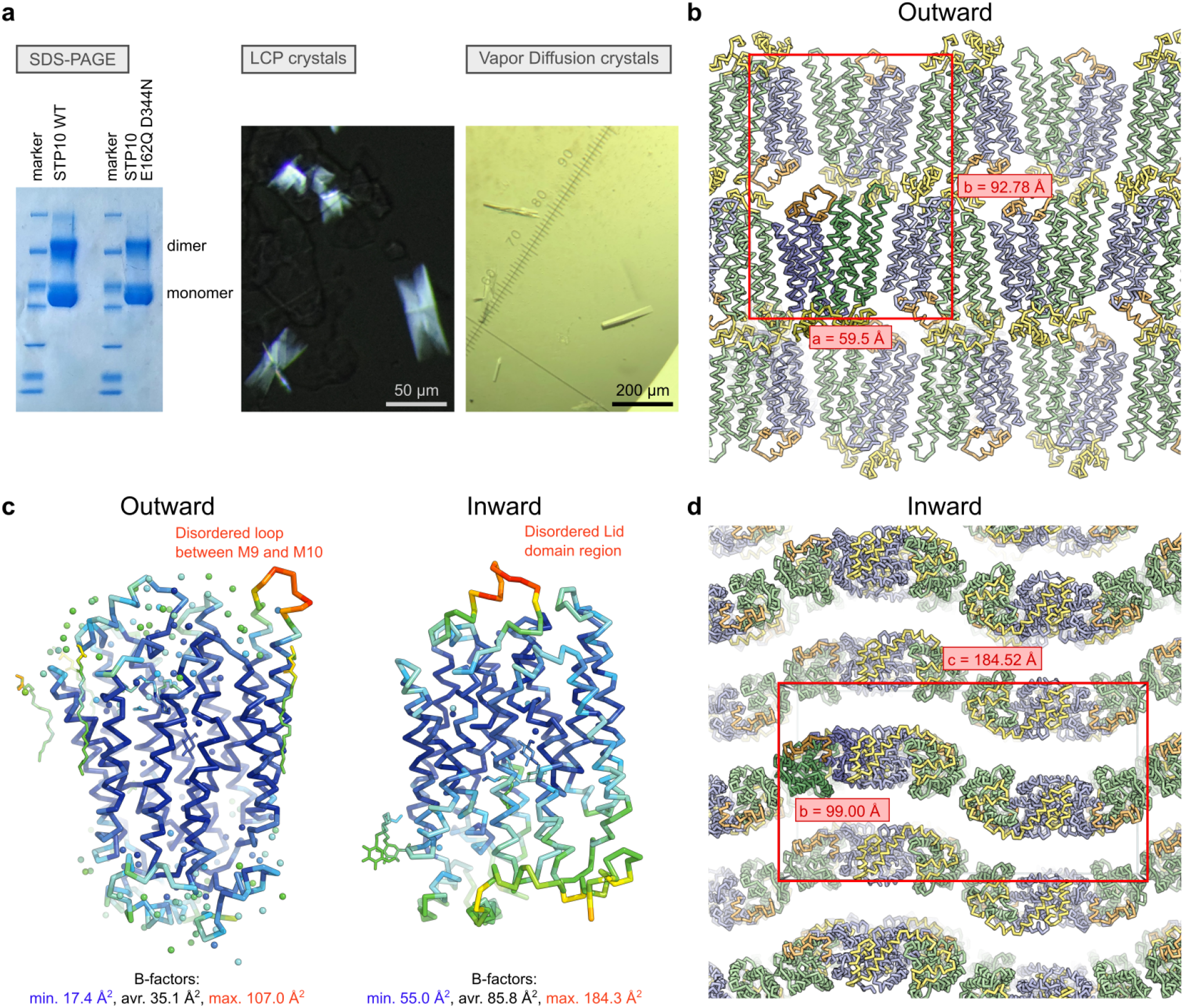
Crystals and components of the asymmetric unit. **a)** SDS-PAGE gel of STP10 protein and polarized light photo of STP10 wild type crystals and light photo of STP10 E162Q D344N crystals. **b)** Asymmetric unit and crystal packing of STP10 wild type. The unit cell is viewed perpendicular to the ab-plane, and the a and b axis highlighted in red. The asymmetric unit contains one molecule of STP10, as highlighted in darker colors. The packing is an example of type I packing normally obtained by LCP crystallography with the transmembrane regions packing in a lipid bilayer and a relatively low solvent content (58%). **c)** The backbone of STP10 outward occluded structure and inward open structure colored by the atomic displacement factor (B-factor) with a rainbow gradient from low/blue to high/red. There is a disordered loop between M9 and M10 with a significantly higher B-factor than the rest of the model in the outward structure and a disordered part of the Lid domain with significant higher B-factor than the rest of the model in the inward open structure. **d)** Asymmetric unit and crystal packing of STP10 E162Q D344N. The unit cell is viewed perpendicular to the bc-plane, and the b and c axis highlighted in red. The asymmetric unit contains one molecule of STP10, as highlighted in darker colors.

**Extended Data Fig. S3.**
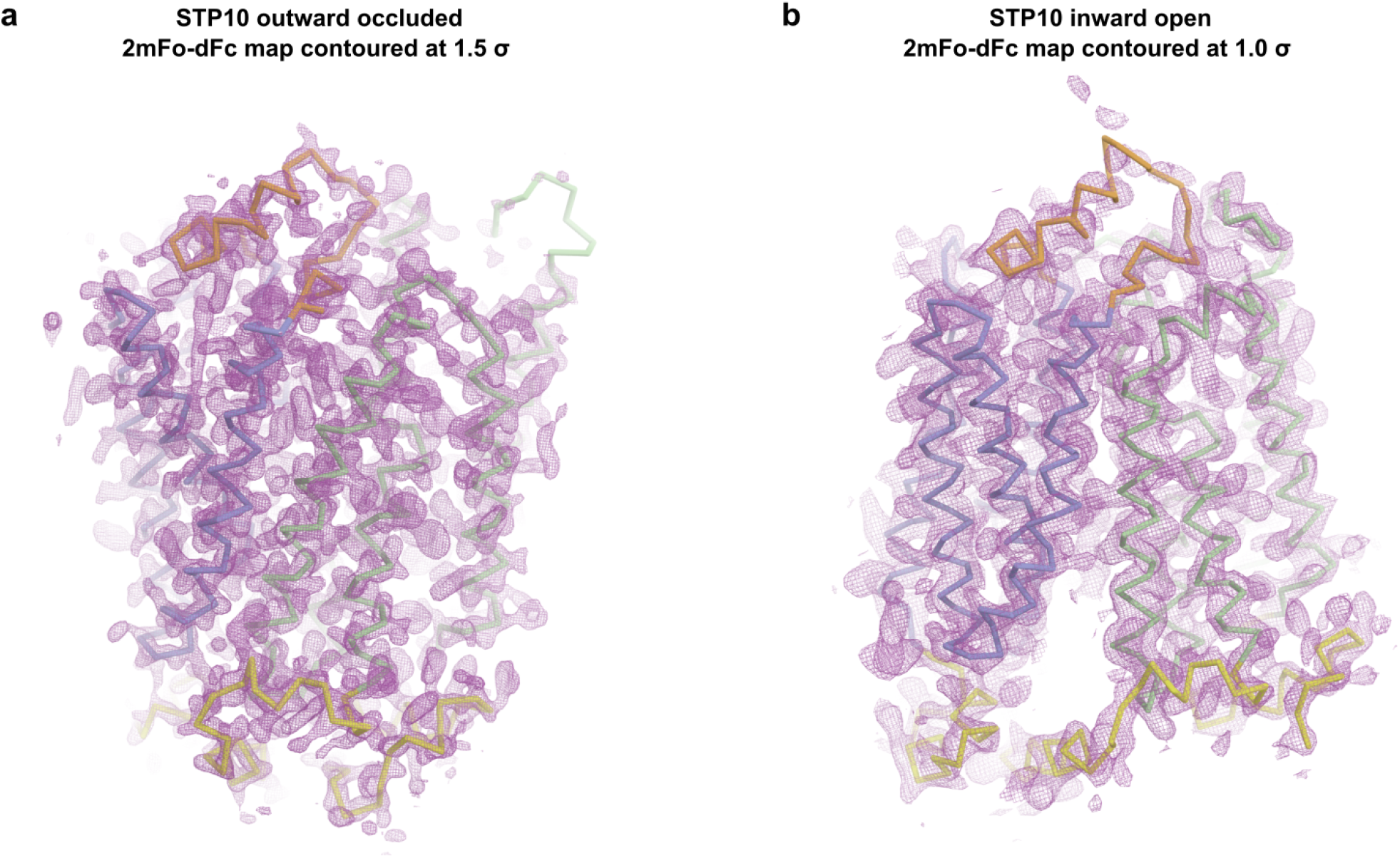
Electron density for the STP10 outward occluded structure and the STP10 inward open structure. **a)** Weighted 2FoFc density at 1.5 sigma of the asymmetric unit of 1.8 Å resolution STP10 outward occluded structure with the final model overlaid. **b)** Weighted 2FoFc density at 1.0 sigma of the asymmetric unit of 2.6 Å resolution STP10 inward open structure with the final model overlaid.

**Extended Data Fig. S4.**
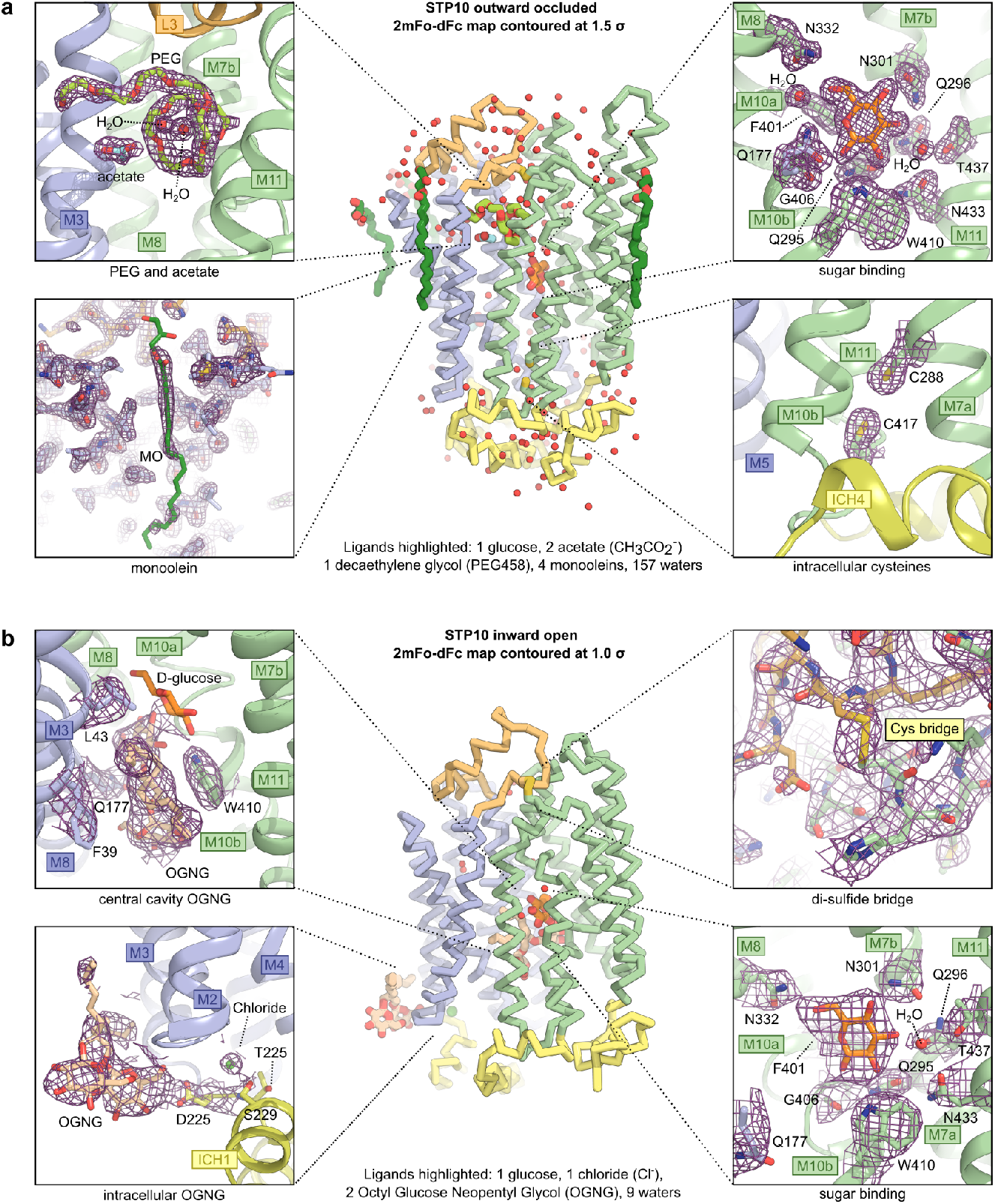
Electron density for selected components of STP10 structures. **a)** Backbone representation of STP10 outward structure with all heterologous molecules found in the density highlighted. Besides STP10 the model contains 1 glucose, 2 acetate, 1 PEG 458, 4 monoolein molecules and 157 waters. The four inserts highlight quality of the electron density displayed by the weighted 2FoFc density at 1.5 sigma for the PEG and acetate, glucose, intracellular cysteines and of the monoolein, which was weaker and is clearer at lower sigma levels than 1.5. **b)** Backbone representation of STP10 inward structure with all the heterologous molecules found in the density highlighted. Besides STP10 the model contains 1 glucose, 1 chloride ion, 2 OGNG and 9 waters. The four inserts highlight quality of the electron density displayed by the weighted 2FoFc density at 1.0 sigma for the two OGNG, glucose and the disulfide bridge.

**Extended Data Fig. S5.**
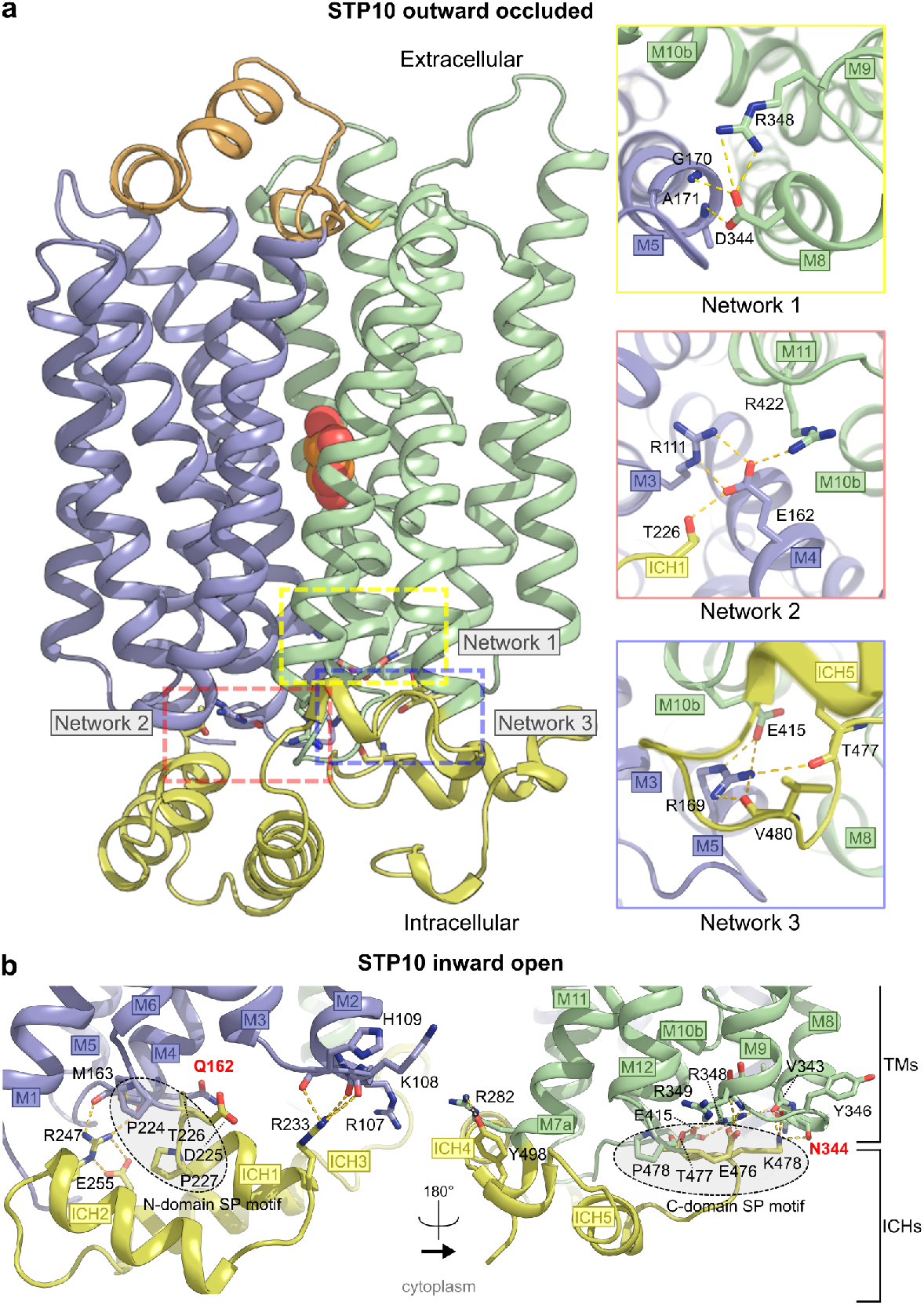
The intracellular gate in the two STP10 structures. **a)** View of the STP10 outward occluded structure perpendicular to the membrane with the three key interdomain salt bridge networks highlighted in colored squares. In particular constituted by the double salt bridge from D344(M8) to the main chain nitrogen of Gly170(M5) and Ala171(M5) (network 1) and the double salt bridge from Glu162 (M4) to Arg422(M11), Thr226 (IC1) and R111(M3) (network 2) as well as and from Arg169(M5) to E415 (M10), the main chain carbonyls of Thr477 (IC5) and Val480 (IC5) (network 3). These regions are perfectly conserved in all STPs (Extended Data Fig. S1) and several bacterial symporters, and have also been observed in human sugar facilitators. **b)** Close-up view of the N domain and C domain at the cytosolic side in the STP10 inward open structure. In the inward open conformation, interactions between the ICH domain and the transmembrane N and C domains are maintained. Interactions between ICH and the two transmembrane domain residues are highlighted by yellow dashes. The mutant residues Q162 and N344 that broke the stabilizing networks are highlighted in red. The positions of the SP motifs are highlighted (dotted eclipses).

**Extended Data Fig. S6.**
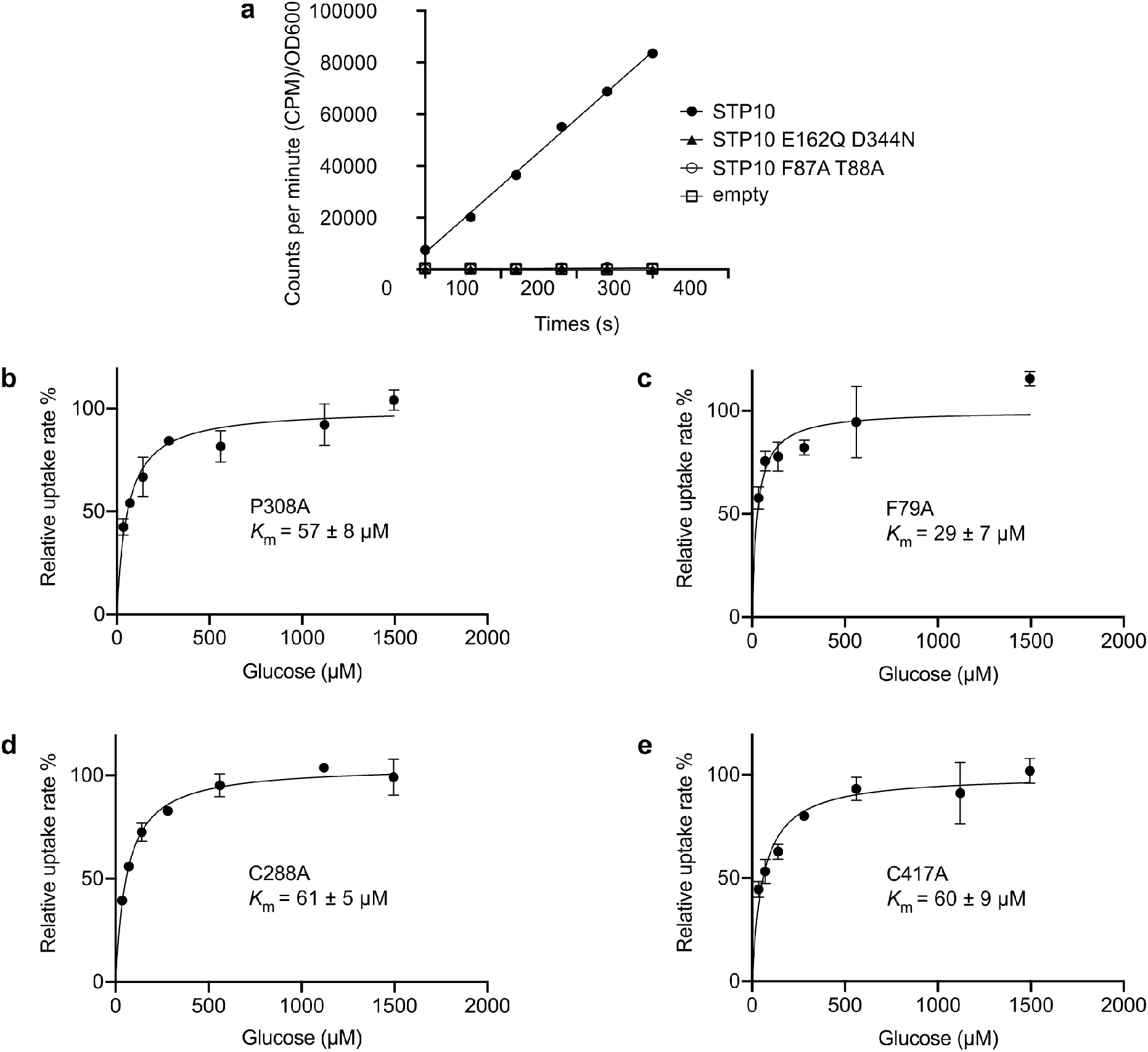
Functional characterization of STP10 mutants. **a)** Uptake of glucose into EBY.VW4000 yeast strain expressing STP10 (black circles), STP10 E162Q D344N (black triangle), STP10 F87A T88A (empty circles) or empty plasmid (empty squares) per OD600 of cells at an initial outside concentration of 100 μM glucose at pH 5.0. **b-d)** Michaelis-Menten fit to glucose titration of STP10 mutants at pH 5.0. Data represents mean ± SD of three or more replicate experiments.

**Extended Data Fig. S7.**
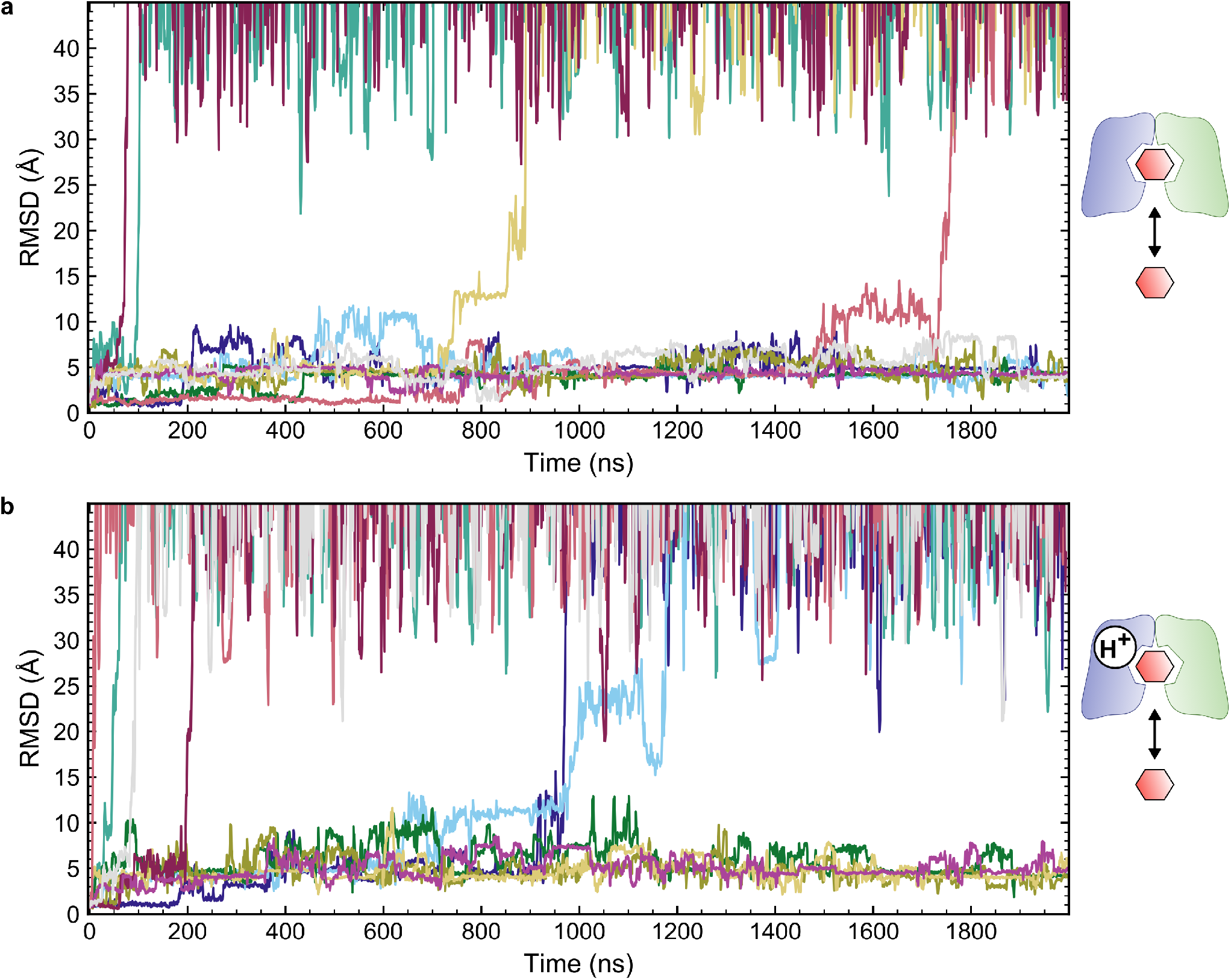
Molecular dynamics simulations of the glucose binding site of STP10 inward open states. **a)** R.m.s.d. ligand plot of the charged inward open state simulations. The glucose leaves the inward state in 4 of 10 independent repeats with Asp42 charged. **b)** R.m.s.d. ligand plot of the neutral inward open state simulations. The glucose leaves the inward state in 6 of 10 independent repeats with Asp42 neutral. Repeats are represented by different color traces same as for Fig. 3d. The ligand RMSD is calculated after aligning the protein structure to the initial model.

**Extended Data Fig. S8.**
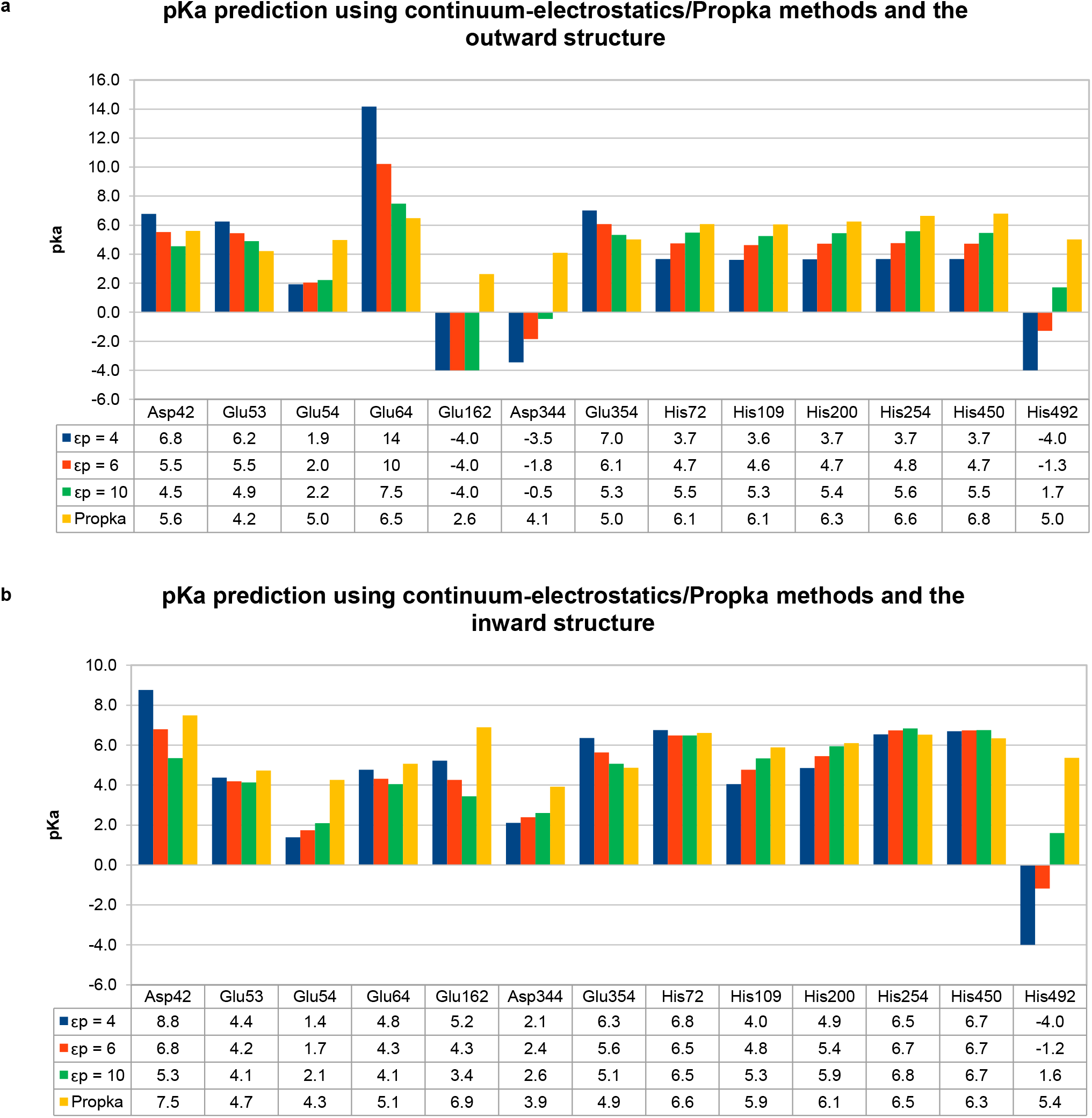
Continuum electrostatics and empirical p*K*_a_ calculations for the two structures. **a)** pK_1/2_ for titratable residues in STP10 for the outward crystal structure with different dielectric constant values for the protein (4, 6 and 10). Glu162 has pK_1/2_ values < −4 for all three dielectric constant values. His492 has pK_1/2_ < −4 for a dielectric constant of 4. Predicted p*K*_a_ using Propka3.0, an empirical method. See subsection Protonation states assignment in Methods for calculation details. **b)** pK_1/2_ for titratable residues in STP10 for the inward crystal structure with different dielectric constant values for the protein (4, 6 and 10). Predicted p*K*_a_ using the empirical method Propka3.0. See subsection Protonation states assignment in Methods for calculation details.

**Extended Data Table S1.** Data collection and refinement statistics.

| Name<br>Type<br>State | STP10<br>native<br>outward occluded | STP10 E162Q/D344N<br>native<br>inward open | STP10 E162Q/D344N<br>bromide<br>inward open |
| --- | --- | --- | --- |
| **Data Collection** |  |  |  |
| Space group | P 21 21 21 | C 2 2 21 | C 2 2 21 |
| Cell dimensions |  |  |  |
| a, b, c (Å) | 59.5 92.8 119.8 | 83.9 99.0 184.5 | 84.2 98.4 185.0 |
| alpha, beta, gamma (deg) | 90 90 90 | 90 90 90 | 90 90 90 |
| Monomers per asym. unit. | 1 | 1 | 1 |
| Wavelength (Å) | 0.9794 | 0.9800 | 0.9203 |
| Number of reflections measured | 676,392 | 152,793 | 243,873 |
| Number of unique reflections | 61,268 | 23,016 | 18,668 |
| Resolution (Å) | 46.39-1.81 (1.84-1.81)<sup>a</sup> | 92.26-2.64 (2.69-2.64)<sup>a</sup> | 24.88-3.50 (4.00-3.50)<sup>a</sup> |
| R<sub>meas</sub> (%) | 17.6 (262.3) | 13.44 (149.6) | 137.9 (103.1) |
| Mean I/σ(I) | 7.7 (1.03) | 6.6 (0.93) | 9.9 (3.03) |
| CC(1/2) | 99.8 (65.1) | 98.7(73.9) | 93.9 (96.8) |
| Completeness (%) | 100 (98.7) | 100 (99.9) | 99.6 (99.9) |
| Redundancy | 11.0 (11.0) | 6.6 (6.8) | 13.0 (13.1) |
| **Refinement** |  |  |  |
| Resolution (Å) | 46.39-1.81 (1.875-1.81) | 19.52-2.64 (2.734-2.64) |  |
| No. reflections (work/free) | 61,084 / 3,030 | 22,480 / 1,092 |  |
| R<sub>work</sub> (%) | 18.74 | 24.66 |  |
| R<sub>free</sub> (%) | 21.20 | 27.79 |  |
| No. of Atoms |  |  |  |
| Protein | 3,771 | 3760 |  |
| Ligands | 151 | 91 |  |
| Waters | 157 | 9 |  |
| Average B Factors (Å<sup>2</sup>) |  |  |  |
| Overall | 37.45 | 85.87 |  |
| Protein | 36.50 | 85.75 |  |
| Ligands | 55.13 | 92.33 |  |
| Waters | 43.21 | 72.35 |  |
| RMSD |  |  |  |
| Bond lengths (Å) | 0.007 | 0.005 |  |
| Bond angles (deg) | 0.77 | 0.83 |  |
| Ramachandran Plot Statistics |  |  |  |
| Favored regions | 99.18 | 96.27 |  |
| Allowed regions | 0.82 | 3.73 |  |
| Disallowed regions | 0.0 | 0.0 |  |
| Deposited model (PDB id) | 7AAQ | 7AAR |  |

<sup>a</sup> Highest resolution shell is shown in parenthesis.

**Extended Data Table S2.** Predicted p*K*_a_ values of Asp42 for outward and inward structures using an ensemble of FEP MD simulations. Free-energy calculations were carried out using the FEP method to analyze the p*K*_a_ shifts of Asp42 with different protonation states of Arg142 and with/without glucose. Five independent repeats were performed for each set of calculations, and the average and standard deviation are reported. ΔG: free energy of charging Asp142 in the protein or in a capped peptide (model). ΔΔG change of free energy of charging Asp142 in the protein relative to a peptide with neutral caps. p*K*_a_ calculated using ΔΔG, T = 310 K, and the experimental p*K*_a_ of Asp in solution (p*K*_a_ = 4.0). ΔG errors are estimated using block averaging with five blocks and then propagated during calculations of ΔΔG and p*K*_a_. See subsection Free-energy perturbation calculations in Methods for detailed information.

| **Outward state with**<br>**glucose and charged Arg142** | $\Delta G$ (kcal/mol) | error (kcal/mol) | $\Delta\Delta G$ (kcal/mol) | error (kcal/mol) | $pK_a$ | error |
| --- | --- | --- | --- | --- | --- | --- |
| repeat 1 | -52 | 0.28 | 7.28 | 0.28 | 9.1 | 0.2 |
| repeat 2 | -53.1 | 0.15 | 6.18 | 0.15 | 8.4 | 0.1 |
| repeat 3 | -52.8 | 0.39 | 6.51 | 0.39 | 8.6 | 0.3 |
| repeat 4 | -53 | 0.1 | 6.33 | 0.11 | 8.5 | 0.08 |
| repeat 5 | -52.5 | 0.22 | 6.79 | 0.22 | 8.8 | 0.2 |
| Average | -52.7 |  | 6.62 |  | 8.7 |  |
| Std. Dev. | 0.43 |  | 0.43 |  | 0.3 |  |
| **Outward state with**<br>**glucose and neutral Arg142** | $\Delta G$ (kcal/mol) | error (kcal/mol) | $\Delta\Delta G$ (kcal/mol) | error (kcal/mol) | $pK_a$ | error |
| repeat 1 | -41.8 | 0.12 | 17.5 | 0.12 | 16 | 0.09 |
| repeat 2 | -41.9 | 0.14 | 17.4 | 0.14 | 16 | 0.1 |
| repeat 3 | -42.1 | 0.31 | 17.2 | 0.31 | 16 | 0.2 |
| repeat 4 | -42.9 | 0.29 | 16.4 | 0.29 | 16 | 0.2 |
| repeat 5 | -41.4 | 0.53 | 17.9 | 0.53 | 17 | 0.4 |
| Average | -42.0 |  | 17.3 |  | 16 |  |
| Std. Dev. | 0.56 |  | 0.56 |  | 0.4 |  |
| **Inward state with**<br>**glucose and charged Arg142** | $\Delta G$ (kcal/mol) | error (kcal/mol) | $\Delta\Delta G$ (kcal/mol) | error (kcal/mol) | $pK_a$ | error |
| repeat 1 | -44.6 | 0.27 | 14.7 | 0.27 | 14 | 0.2 |
| repeat 2 | -47.1 | 0.35 | 12.2 | 0.35 | 13 | 0.2 |
| repeat 3 | -43.4 | 0.75 | 15.9 | 0.75 | 15 | 0.5 |
| repeat 4 | -43.1 | 0.58 | 16.2 | 0.58 | 15 | 0.4 |
| repeat 5 | -45.2 | 0.29 | 14.1 | 0.29 | 14 | 0.2 |
| Average | -44.7 |  | 14.6 |  | 14 |  |
| Std. Dev. | 1.6 |  | 1.6 |  | 1 |  |
| **Inward state with**<br>**glucose and neutral Arg142** | $\Delta G$ (kcal/mol) | error (kcal/mol) | $\Delta\Delta G$ (kcal/mol) | error (kcal/mol) | $pK_a$ | error |
| repeat 1 | -31.8 | 0.74 | 27.5 | 0.74 | 23 | 0.5 |
| repeat 2 | -41.6 | 0.11 | 17.7 | 0.11 | 16 | 0.08 |
| repeat 3 | -35.9 | 0.51 | 23.4 | 0.51 | 21 | 0.4 |
| repeat 4 | -37.8 | 0.17 | 21.5 | 0.17 | 19 | 0.1 |
| repeat 5 | -39.4 | 0.049 | 19.9 | 0.059 | 18 | 0.04 |
| Average | -37.3 |  | 22.0 |  | 19 |  |
| Std. Dev. | 3.7 |  | 3.7 |  | 3 |  |
| **Inward state without**<br>**glucose and charged Arg142** | $\Delta G$ (kcal/mol) | error (kcal/mol) | $\Delta\Delta G$ (kcal/mol) | error (kcal/mol) | $pK_a$ | error |
| repeat 1 | -49.8 | 0.093 | 9.53 | 0.098 | 11 | 0.07 |
| repeat 2 | -46.5 | 0.48 | 12.8 | 0.48 | 13 | 0.3 |
| repeat 3 | -49.3 | 0.33 | 9.95 | 0.34 | 11 | 0.2 |
| repeat 4 | -45.5 | 0.76 | 13.8 | 0.76 | 14 | 0.5 |
| repeat 5 | -48.1 | 0.43 | 11.2 | 0.43 | 12 | 0.3 |
| Average | -47.8 |  | 11.5 |  | 12 |  |
| Std. Dev. | 1.8 |  | 1.8 |  | 1 |  |
| **Model peptide** | $\Delta G$ (kcal/mol) | error (kcal/mol) | | | | |
| Asp | -59.3 | 0.033 |  |  |  |  |

